## Supplementary Figures for "A central protein complex essential for Invasion in *Toxoplasma gondii*"

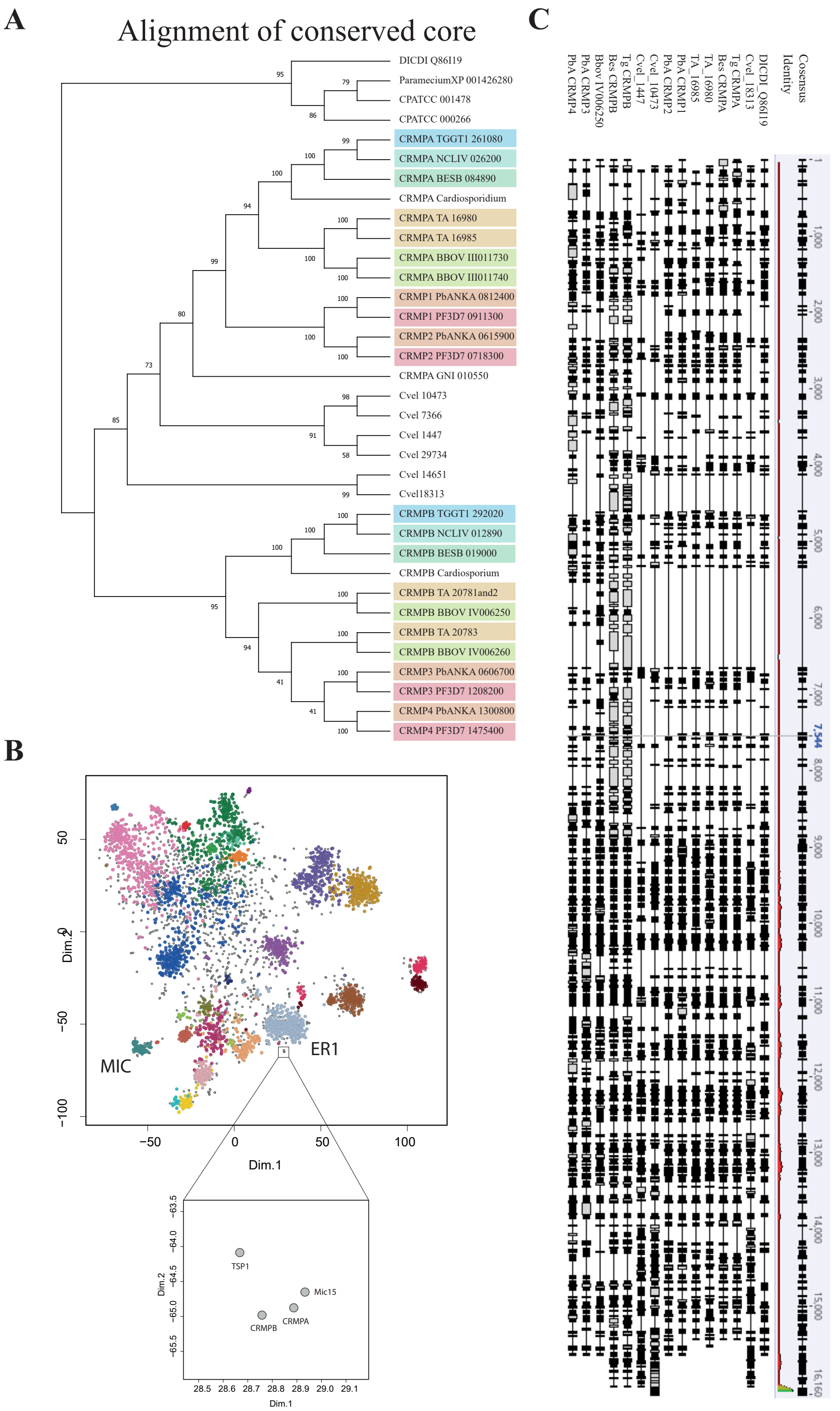
A
Alignment of conserved core
DICDI Q86I19
ParameciumXP 001426280
CPATCC 001478
CPATCC 000266
CRMPA TGGT1 261080
CRMPA NCLIV 026200
CRMPA BESB 084890
CRMPA Cardiosporidium
CRMPA TA 16980
CRMPA TA 16985
CRMPA BBOV III011730
CRMPA BBOV III011740
CRMP1 PbANKA 0812400
CRMP1 PF3D7 0911300
CRMP2 PbANKA 0615900
CRMP2 PF3D7 0718300
CRMPA GNI 010550
Cvel 10473
Cvel 7366
Cvel 1447
Cvel 29734
Cvel 14651
Cvel18313
CRMPB TGGT1 292020
CRMPB NCLIV 012890
CRMPB BESB 019000
CRMPB Cardiosporium
CRMPB TA 20781and2
CRMPB BBOV IV006250
CRMPB TA 20783
CRMPB BBOV IV006260
CRMP3 PbANKA 0606700
CRMP3 PF3D7 1208200
CRMP4 PbANKA 1300800
CRMP4 PF3D7 1475400
B
MIC
ER1
Dim.1
Dim.2
TSP1
Mic15
CRMPB
CRMPA
C
Cosensus
Identity
DICDI_Q86I19
Cvel_18313
Tg CRMPA
Bes CRMPA
TA_16980
TA_16985
PbA CRMP1
PbA CRMP2
Cvel_10473
Cvel_1447
Tg CRMPB
Bes CRMPB
Bbov IV006250
PbA CRMP3
PbA CRMP4

**Figure S1: Phylogenetic analysis and Lopit map placement. A** Phylogenetic relationship of CRPMs is shown after alignment of the conserved core region using the PRANK algorithm, bootstrap analysis (100x) supports most branches. Compare with Figure 1 A. **B** Spatial proteomic map of all *T. gondii* proteins[30], the micronemal cluster (MIC) and the ER1 cluster are indicated. Insert on the right below the ER cluster shows a separate "minicluster" of CRMPB and CRMPA, grouping together with Thrombospondin typ 1 domain-containing protein (TSP1; TGGT1_277910) and Mic15 (TGGT1_247195). **C** A local conservation map and local protein identity and conservation. Insertion are indicated by grew boxes.

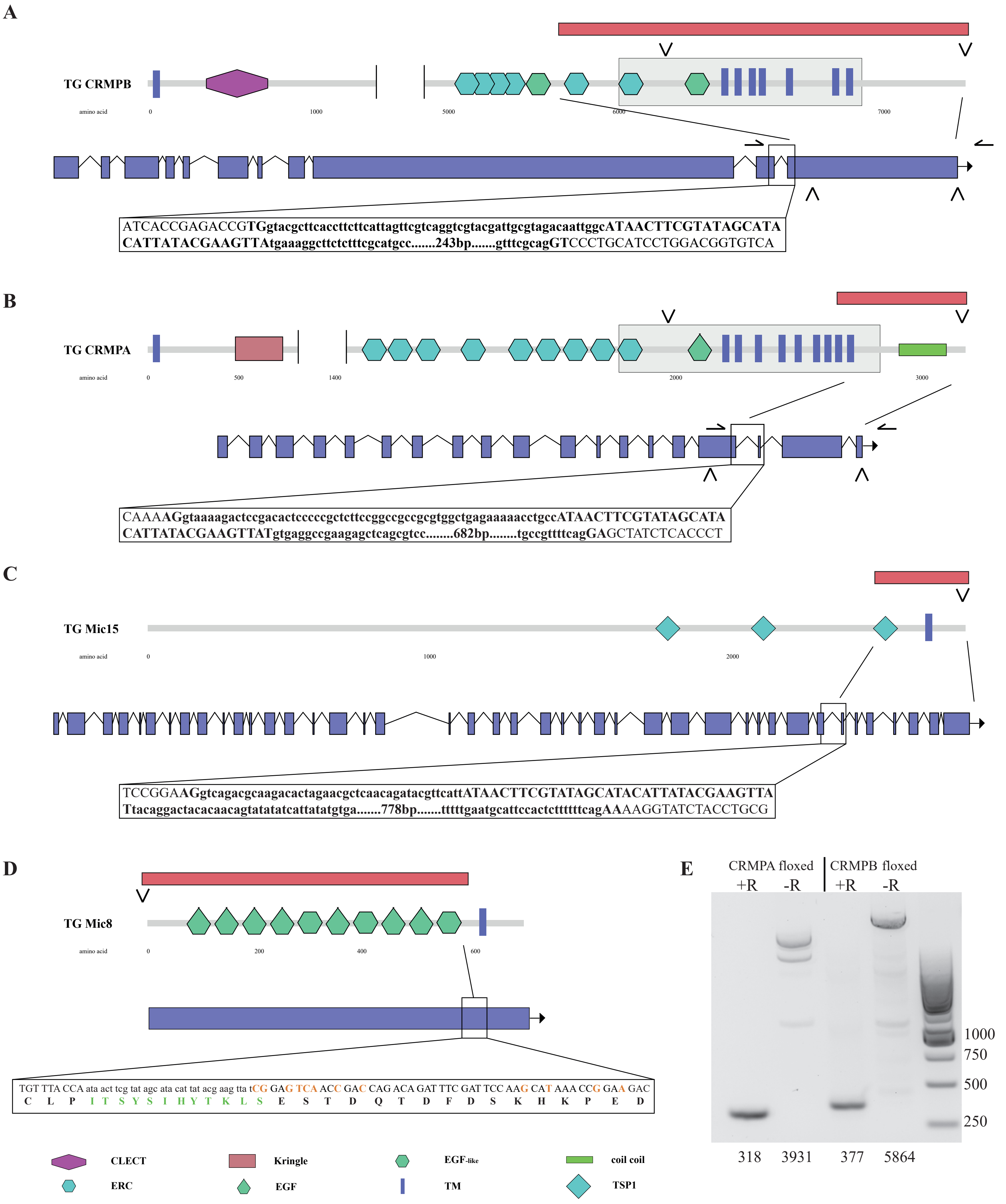

A
TG CRMPB
amino acid
0
1000
5000
6000
7000
ATCACCGAGACCGTGgtacgcttcaccttcttcattagttcgtcaggtcgtacgattgcgtagacaattggcATAACTTCGTATAGCATA
CATTATACGAAGTTAtgaaaggcttctctttcgcatgcc.......243bp.......gtttcgcagGTCCCTGCATCCTGGACGGTGTCA
B
TG CRMPA
amino acid
0
500
1400
2000
3000
CAAAAGgtaaaagactccgacactcccccgctcttccggccgccgcgtggctgagaaaaacctgccATAACTTCGTATAGCATA
CATTATACGAAGTTATgtgaggccgaagagctcagcgtcc........682bp........tgccgttttcagGAGCTATCTCACCCT
C
TG Mic15
amino acid
0
1000
2000
TCCGGAAGgtcagacgcaagacactagaacgctcaacagatacgttcattATAACTTCGTATAGCATACATTATACGAAGTTA
Ttacaggactacacaacagtatatatcattatatgtga.......778bp.......tttttgaatgcattccactcttttttcagAAAAGGTATCTACCTGCG
D
TG Mic8
amino acid
0
200
400
600
TGT TTA CCA ata act tcg tat agc ata cat tat acg aag tta tCG GAG TCA ACC GAC CAG ACA GAT TTC GAT TCC AAG CAT AAA CCG GAA GAC
C L P I T S Y S I H Y T K L S E S T D Q T D F D S K H K P E D
E
CRMPA floxed
CRMPB floxed
+R
-R
+R
-R
1000
750
500
250
318
3931
377
5864
CLECT
Kringle
EGF-like
coil coil
ERC
EGF
TM
TSP1

**Figure S2: Gene tagging strategies. A** Shown is an overview of the domain architecture and the gene CRMPB of *T. gondii*. Arrowheads indicate the position of the C-terminal (including the C-terminal LoxP site) and extracellular tag in the gene and the protein model. Position of the LoxP in the intron and the local sequence is shown, relative position within the domain model is indicated by a line and lost sequence after rapamycin induction is indicated by a red bar. **B** Overview of the domain architecture and the gene of CRMPA. Note that only the last transmembrane domain and the C-terminus is lost upon induction with rapamycin. **C** Overview of the domain architecture and the gene Mic15. Position of the LoxP within the intron and the part of the gene lost after rapamycin induction is indicated. **D** Overview of domain architecture of Mic8. The upstream LoxP sequence is 6 bp upstream of the start site (position indicated by an arrow head), the internal LoxP site is 3 bp after the last EGF-like domain inserted as coding as indicated, inserting the protein sequence ITSYSIHYTKLS into the protein (coding sequence in lower case, protein sequence in green). Shielding mutations inserted are indicated with orange letters. Several domains are indicated: CLECT (c-type Lectin or carbohydrate recognition domain), ERC (Ephrin-receptor like), EGF-like (Epidermal growth factor-like), TM (transmembrane domain), Kringle (Kringle domain), EGF (Epidermal growth factor), coil coil (alpha helical coil coil domains), TSP1 (thrombospondin type-1 repeat). **E** Genotyping of RH-CRMPA-3HA$^{floxed}$ and RH-CRMPB-3HA$^{floxed}$ after 72 h induction with rapamycin. The respective primers bind outside of the floxed region and are indicated in **A** and **B** with arrows. Estimated sizes are indicated below in basepairs.

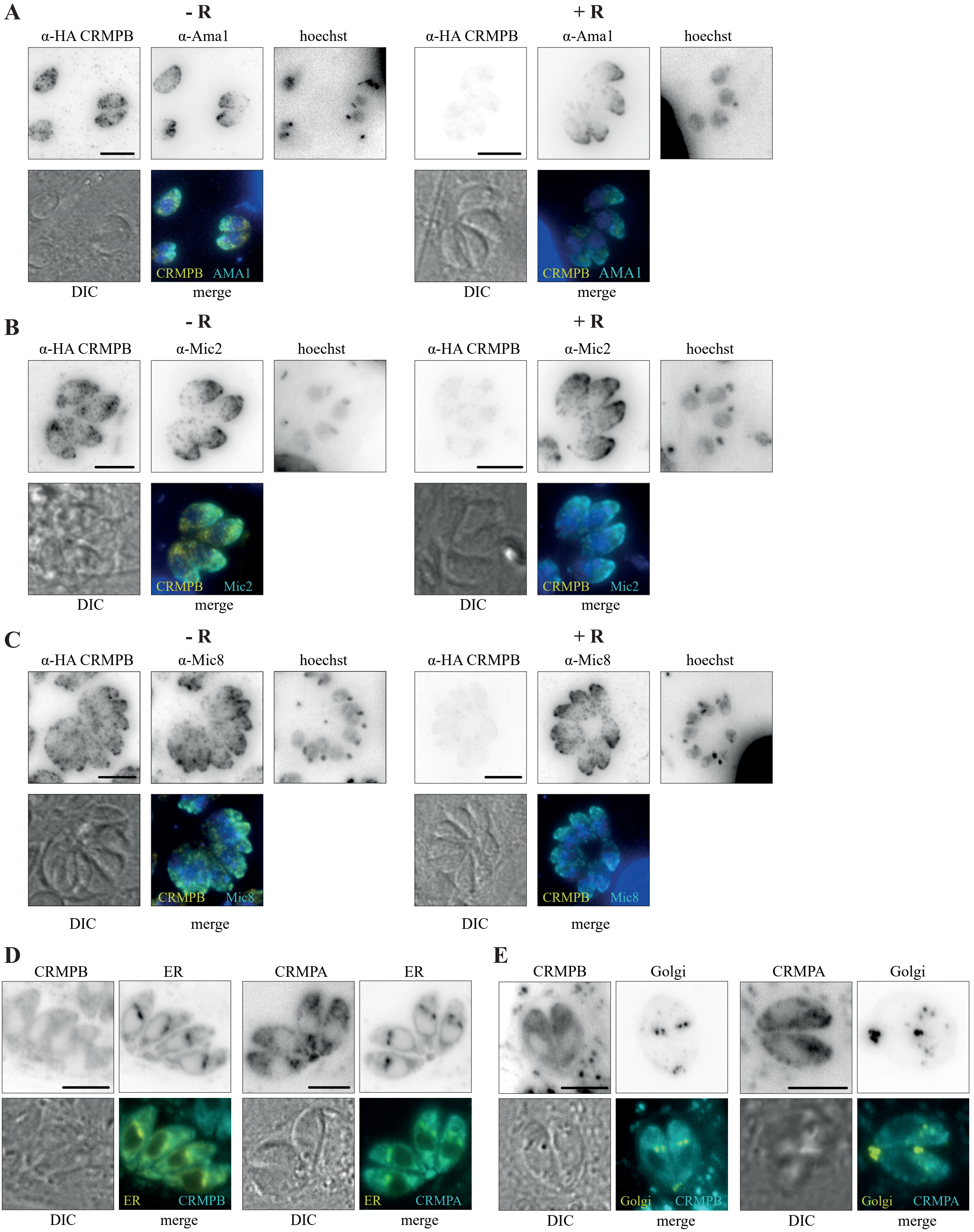

A
- R
+ R
α-HA CRMPB
α-Ama1
hoechst
α-HA CRMPB
α-Ama1
hoechst
CRMPB AMA1
DIC
merge
CRMPB AMA1
DIC
merge
B
- R
+ R
α-HA CRMPB
α-Mic2
hoechst
α-HA CRMPB
α-Mic2
hoechst
CRMPB Mic2
DIC
merge
CRMPB Mic2
DIC
merge
C
- R
+ R
α-HA CRMPB
α-Mic8
hoechst
α-HA CRMPB
α-Mic8
hoechst
CRMPB Mic8
DIC
merge
CRMPB Mic8
DIC
merge
D
CRMPB
ER
CRMPA
ER
ER CRMPB
ER CRMPA
DIC
merge
DIC
merge
E
CRMPB
Golgi
CRMPA
Golgi
Golgi CRMPB
Golgi CRMPA
DIC
merge
DIC
merge

**Figure S3: Colocalization of CRMP and micronemal proteins.** The abundance and localization of **A** Ama1, **B** Mic2 and **C** Mic8 relative to CRMPB was analyzed in presence and absence (72 h rapamycin) of CRMPB in RH-CRMPB-3HA$^{floxed}$. **D** Parasites with extracellular SNAP tagged CRMPB (RH-SNAP-CRMPB-3HA$^{floxed}$ CRMPA-sYFP2) or extracellular Halo tagged CRMPA (RH-Halo-CRMPA-3HA$^{floxed}$ CRMPB-sYFP2) were transiently transfected with **D** ERD-GFP and **E** GalNac YFP (TGGT1_259530-YFP)[68] and labelled (SNAP-Cell or Halo-Janelia 646) and imaged live 48 h post transfection. Scale bars are 5 µm.

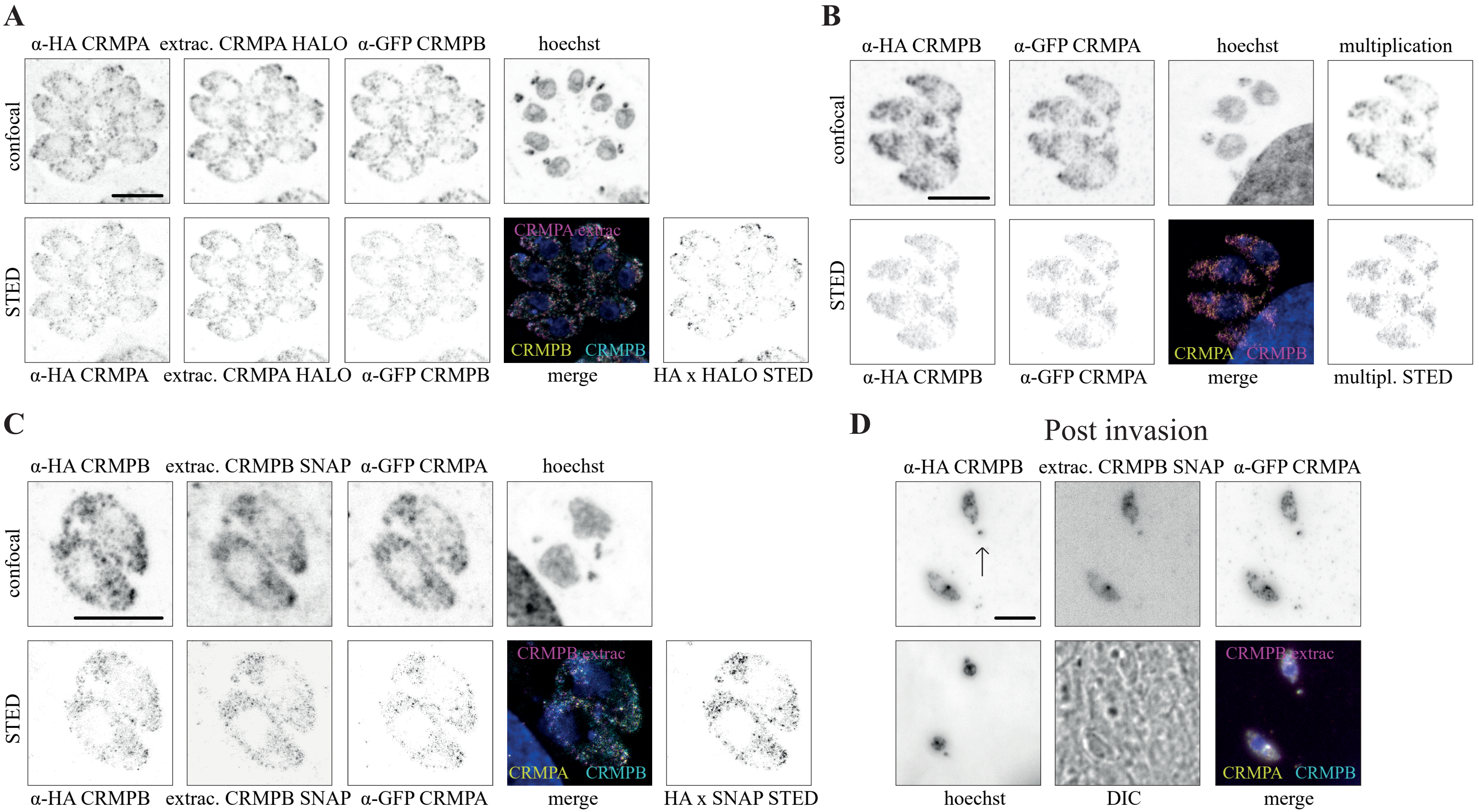
A
α-HA CRMPA
extrac. CRMPA HALO
α-GFP CRMPB
hoechst
confocal
STED
CRMPA extrac
CRMPB
CRMPB
α-HA CRMPA
extrac. CRMPA HALO
α-GFP CRMPB
merge
HA x HALO STED
B
α-HA CRMPB
α-GFP CRMPA
hoechst
multiplication
confocal
STED
CRMPA
CRMPB
α-HA CRMPB
α-GFP CRMPA
merge
multipl. STED
C
α-HA CRMPB
extrac. CRMPB SNAP
α-GFP CRMPA
hoechst
confocal
STED
CRMPB extrac
CRMPA
CRMPB
α-HA CRMPB
extrac. CRMPB SNAP
α-GFP CRMPA
merge
HA x SNAP STED
D
Post invasion
α-HA CRMPB
extrac. CRMPB SNAP
α-GFP CRMPA
CRMPB extrac
CRMPA
CRMPB
hoechst
DIC
merge

**Figure S4: Extracellular tagging of CRMPs. A** Localization of HALO (Janelia fluor 646) within the extracellular domain of CRMPA (see Figure S2) in respect to C-terminally tagged CRMPA (3HA) and C-terminally tagged CRMPB (sYFP2). **B** Localization of RH-CRMPB-3HA$^{floxed}$ CRMPA-sYFP2 was stained with for HA (Star635P) and sYFP2 (Atto594). Compare with Figure 2 A. **C** Localization of SNAP (Sir647) within the extracellular domain of CRMPB (see Figure S2) in respect to C-terminally tagged CRMPB (3HA) and C-terminally tagged CRMPA (sYFP2). **D** Parasites with extracellular CRMPB SNAP (Janelia 646) after pulse invasion in the presence of proponol. The putative invasion site is marked with an arrow. Scale bars are 5 μm.

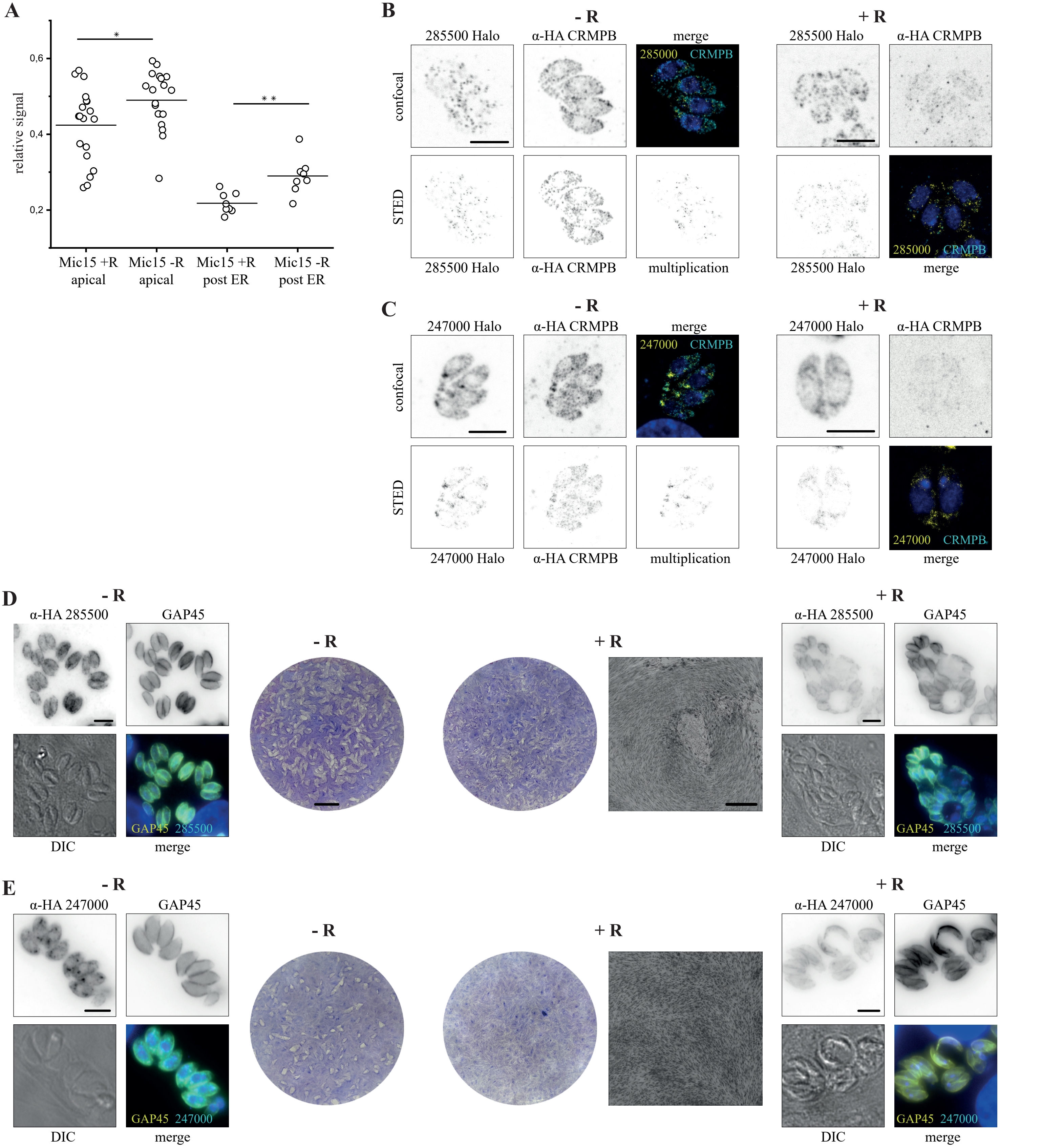
A
B
C
D
E
relative signal
0,6
0,4
0,2
*
* *
Mic15 +R apical
Mic15 -R apical
Mic15 +R post ER
Mic15 -R post ER
- R
+ R
285500 Halo
α-HA CRMPB
merge
confocal
STED
285000 CRMPB
multiplication
247000 Halo
247000 CRMPB
α-HA 285500
GAP45
DIC
GAP45 285500
α-HA 247000
GAP45 247000

**Figure S5: Characterization of potential CRMP interactors.** **A** The relative localization of Mic15 was quantified in the presence and absence of CRMPB. Analysis was performed manually (apical of total fluorescence) and automatically (post ER of total fluorescence). For details see methods section. **B** Colocalization of TGGT1_285500 (Halo) with CRMPB and localization of 285500 in the absence of CRMPB (72h). **C** Colocalization of the tetratricopeptide repeat containing protein (TGGT1_247000) (Halo) with CRMPB and localization of 247000 in the absence of CRMPB (72h). **D** Localization of 3HA tagged 285500 + / - rapamycin (72 h). Plaque assay after 7 days shows a mild growth phenotype. **E** Localization of 3HA tagged 247000 + / - rapamycin (72 h). No plaques could be detected 7 days after rapamycin induction. Note that there is some crosstalk with the Gap45 signal in C and D. Scale bars of the plaque assay are 5 mm (scale bar of the higher resolution area is 500 μm) all other scale bars are 5 μm.

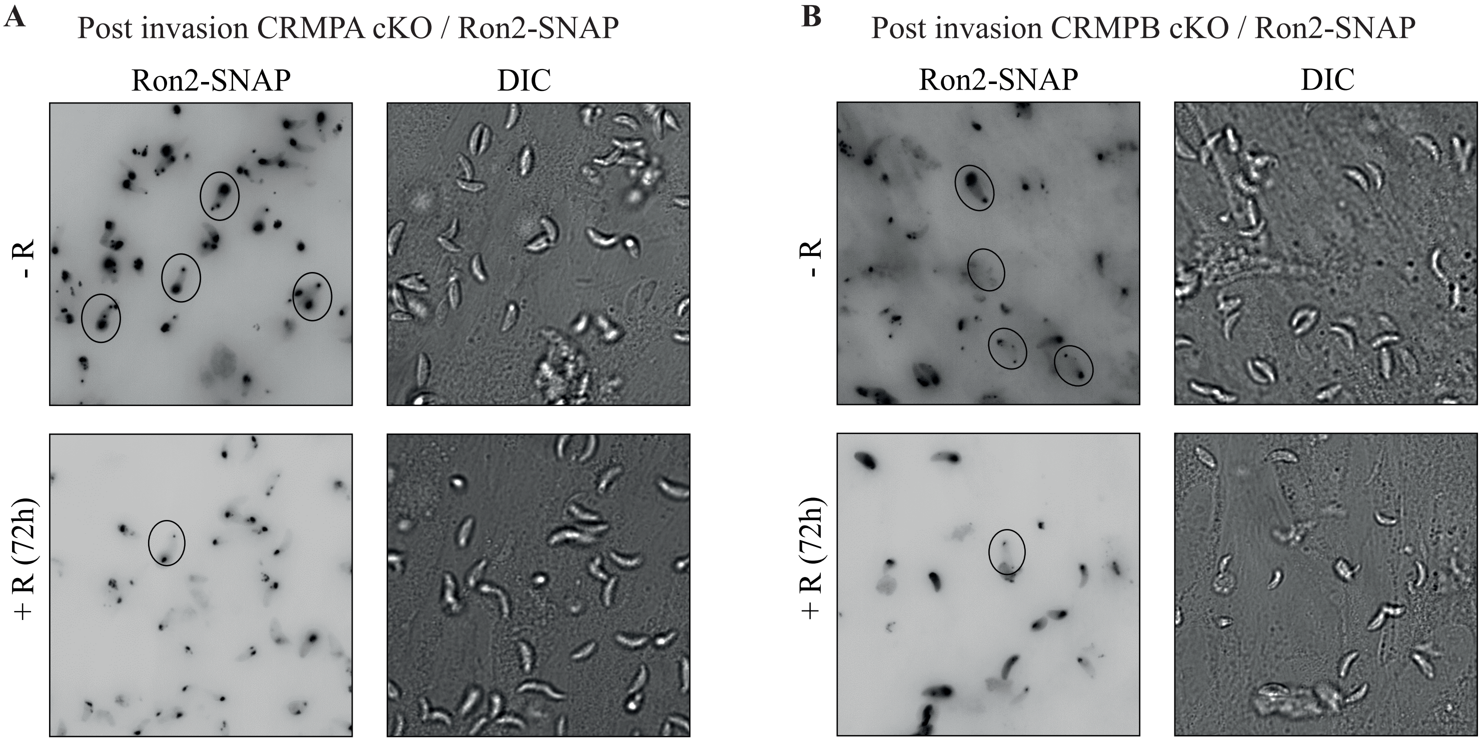
A
Post invasion CRMPA cKO / Ron2-SNAP
Ron2-SNAP
DIC
- R
+ R (72h)
B
Post invasion CRMPB cKO / Ron2-SNAP
Ron2-SNAP
DIC
- R
+ R (72h)

**Figure S6: Analysis of invasion using Ron2-SNAP.** In the cKO parasites of CRMPB and CRMPA, Ron2 was endogenously tagged with SNAP. Shown are maximum projections after 20 minutes of invasion (72 h post rapamycin induction), invaded parasites are circled. Both invaded parasites in the plus rapamycin group represent the only tachyzoite with a clear “invaded” Ron2 staining pattern, minus rapamycin are representative stacks.

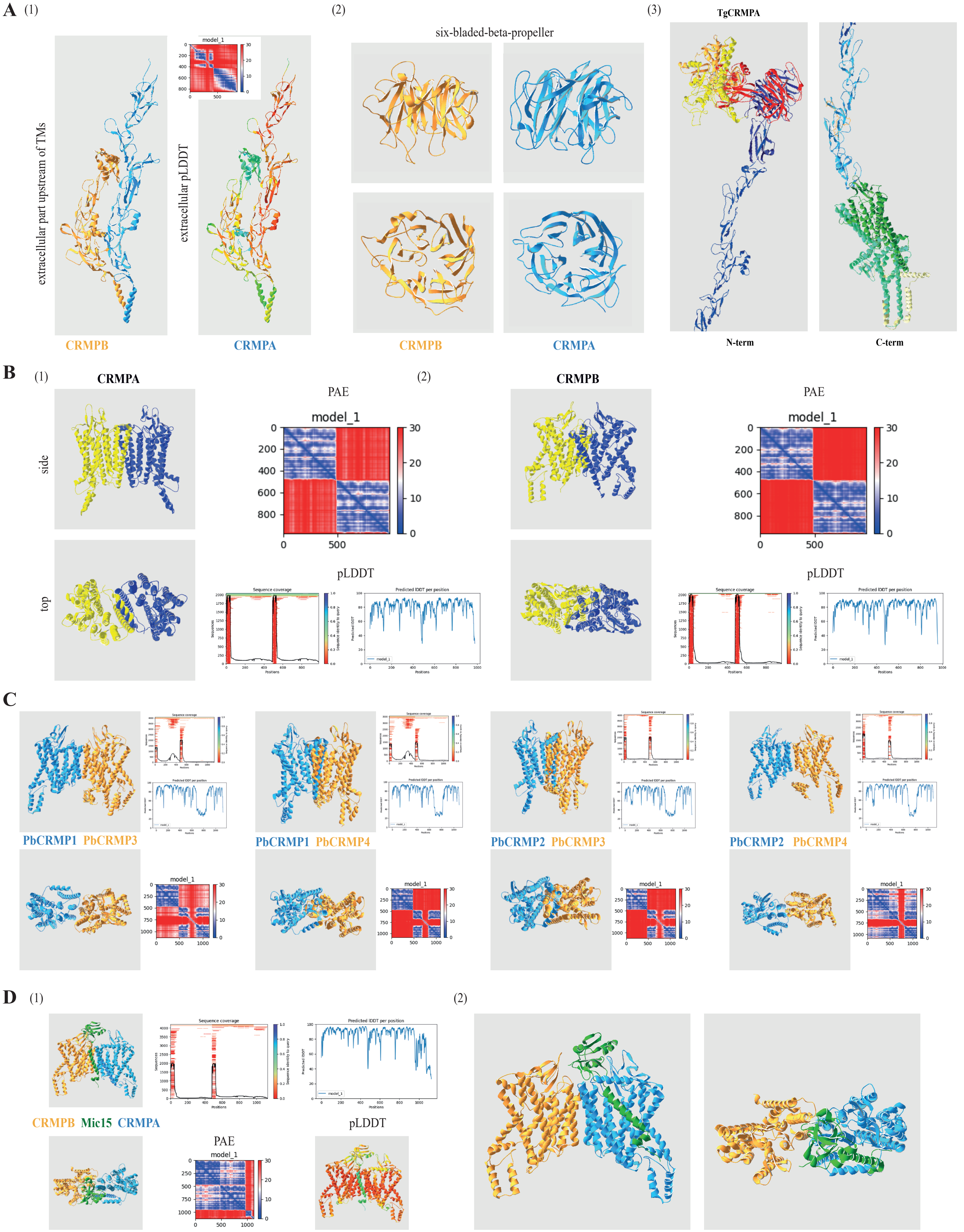
A
(1)
extracellular part upstream of TMs
extracellular pLDDT
model_1
CRMPB
CRMPA
(2)
six-bladed-beta-propeller
CRMPB
CRMPA
(3)
TgCRMPA
N-term
C-term
B
(1)
CRMPA
side
top
PAE
model_1
pLDDT
Sequence coverage
Predicted lDDT per position
(2)
CRMPB
PAE
model_1
pLDDT
C
PbCRMP1 PbCRMP3
PbCRMP1 PbCRMP4
PbCRMP2 PbCRMP3
PbCRMP2 PbCRMP4
model_1
D
(1)
Sequence coverage
Predicted lDDT per position
CRMPB Mic15 CRMPA
PAE
model_1
pLDDT
(2)

**Figure S7: Structural modeling of CRMPs. A** Multimer homodimer structure prediction of (1) the extracellular part directly upstream of the TMs of TgCRMPA/B. (2) The six-bladed-beta-propeller was found in the extracellular domain of both TgCRMPA and TgCRMPB. (3) Structure of TgCRMPA modeled with RobettaFold, Modeling was performed of 9 fragments, with 60 amino acid overlap. Each fragment is shown in a separate color. **B** Homodimer modeling for (1) CRMPA and (2) CRMPB resulted in overlay of proteins. The PAE scores showed good relative prediction for single chains, but indicates that both chains do not interact, despite the high IDDT values. Sequence alignment blots indicate the number of sequences that could be aligned to each position. **C** Multimer heterodimer prediction of PbCRMP1/3, PbCRMP1/4, PbCRMP2/3 and PbCRMP2/4. CRMPA clade members are colored light blue, CRMPB clade members are colored orange. PAE scores indicate that PbCRMP1/3 and PbCRMP2/4 do interact and PbCRMP1/4 and PbCRMP2/3 do not interact. PbCRMP3 and PbCRMP4 are predicted in two fragments with an unstructured part (indicated by the low local pLDDT score).
