## Supplementary data for "A central protein complex essential for Invasion in *Toxoplasma gondii*"

List of gRNAs

| Name | GeneID | gRNA | location |
|---|---|---|---|
| CRMPA | TGGT1_261080 | **G**TCCTGCTTACAGACCGTCGT | C-term |
| CRMPA | TGGT1_261080 | GAGCTCTTCGGCCTCACGGC | intron |
| CRMPA | TGGT1_261080 | **G**CGGCTTGCTCGAGGAAGCAG | extracellular tag |
| CRMPB | TGGT1_292020 | **G**TCTCTAAGCTACTCCTGTTT | C-term |
| CRMPB | TGGT1_292020 | **G**CAGCTAGTCTCGAGTTGTTC | 1089 bp before ATG |
| CRMPB | TGGT1_292020 | **G**TTGCGTAGACAATTGGCGAA | intron |
| CRMPB | TGGT1_292020 | GCGCGTTACAGATGCGATCG | extracellular tag |
| Mic15 | TGGT1_247195 | GCTGGACACGTCCACCCAAG | C-term |
| Mic15 | TGGT1_247195 | **G**CAACAGATACGTTCATTTAC | intron |
| TSP1 | TGGT1_277910 | **G**CCGTGACACCCGAATCCTTT | C-term |
| TSP1 | TGGT1_277910 | GAAGAAGAACGGAAAATCTC | pre ATG |
| TRP | TGGT1_247000 | **G**TCTCCTTCACTGACTTGCTG | C-term |
| TRP | TGGT1_247000 | **G**CGTCACAAATCGAACAGCCA | pre ATG |
| hypothetical | TGGT1_285500 | **G**CATGTGATGATTGTGAAAAG | C-term |
| hypothetical | TGGT1_285500 | **G**TTCTGGATAGCCCATTCGCG | pre ATG |
| Ron2 | TGGT1_300100 | **G**AAACGTCCGAAAAACAGACG | C-term |
| Mic8 | TGGT1_245490 | **G**TTGGCCTTCATTCTGTCTTG | pre ATG |
| Mic8 | TGGT1_245490 | **G**TCTGTCTGATCAGTGCTTTC | internal after EGF |

Sequence of tags

3xHA

GCTAAAATTGGAAGTGGAGGATACCCGTACGACGTCCCGGACTACGCTGGCTATCCCTATGATGTG
CCCGATTATGCGTATCCTTACGATGTTCCAGATTATGCCTAAATAACTTCGTATAGCATACATTATA
CGAAGTTAT

sYFP2

GCTAAAATTGGAAGTGGAGGACGGATGGTGAGCAAGGGCGAGGAGCTGTTCACCGGGGTGGTGCC
CATCCTGGTCGAGCTGGACGGCGACGTAAACGGCCACAAGTTCAGCGTGTCCGGCGAGGGCGAGG
GCGATGCCACCTACGGCAAGCTGACCCTGAAGCTGATCTGCACCACCGGCAAGCTGCCCGTGCCCT
GGCCCACCCTCGTGACCACCCTGGGCTACGGCGTGCAGTGCTTCGCCCGCTACCCCGACCACATGA
AGCAGCACGACTTCTTCAAGTCCGCCATGCCCGAAGGCTACGTCCAGGAGCGCACCATCTTCTTCA
AGGACGACGGCAACTACAAGACCCGCGCCGAGGTGAAGTTCGAGGGCGACACCCTGGTGAACCGC
ATCGAGCTGAAGGGCATCGACTTCAAGGAGGACGGCAACATCCTGGGGCACAAGCTGGAGTACAA

CTACAACAGCCACAACGTCTATATCACCGCCGACAAGCAGAAGAACGGCATCAAGGCCAACTTCA
AGATCCGCCACAACATCGAGGACGGCGGCGTGCAGCTCGCCGACCACTACCAGCAGAACACCCCC
ATCGGCGACGGCCCCGTGCTGCTGCCCGACAACCACTACCTGAGCTACCAGTCCAAGCTGAGCAAA
GACCCCAACGAGAAGCGCGATCACATGGTCCTGCTGGAGTTCGTGACCGCCGCCGGGATCACTCTC
GGCATGGACGAGCTGTACAAGTAGATAACTTCGTATAGCATACATTATACGAAGTTAT

HALO

GCTAAAATTGGAAGTGGAGGACGGCCTAGGCTCGAGCCAACCACTGAGGATCTGTACTTTCAGAG
CGATAACGATGGATCCGAAATCGGTACTGGCTTTCCATTCGACCCCCATTATGTGGAAGTCCTGGG
CGAGCGCATGCACTACGTCGATGTTGGTCCGCGCGATGGCACCCCTGTGCTGTTCCTGCACGGTAA
CCCGACCTCCTCCTACGTGTGGCGCAACATCATCCCGCATGTTGCACCGACCCATCGCTGCATTGCT
CCAGACCTGATCGGTATGGGCAAATCCGACAAACCAGACCTGGGTTATTTCTTCGACGACCACGTC
CGCTTCATGGATGCCTTCATCGAAGCCCTGGGTCTGGAAGAGGTCGTCCTGGTCATTCACGACTGG
GGCTCCGCTCTGGGTTTCCACTGGGCCAAGCGCAATCCAGAGCGCGTCAAAGGTATTGCATTTATG
GAGTTCATCCGCCCTATCCCGACCTGGGACGAATGGCCAGAATTTGCCCGCGAGACCTTCCAGGCC
TTCCGCACCACCGACGTCGGCCGCAAGCTGATCATCGATCAGAACGTTTTTATCGAGGGTACGCTG
CCGATGGGTGTCGTCCGCCCGCTGACTGAAGTCGAGATGGACCATTACCGCGAGCCGTTCCTGAAT
CCTGTTGACCGCGAGCCACTGTGGCGCTTCCCAAACGAGCTGCCAATCGCCGGTGAGCCAGCGAAC
ATCGTCGCGCTGGTCGAAGAATACATGGACTGGCTGCACCAGTCCCCTGTCCCGAAGCTGCTGTTC
TGGGGCACCCCAGGCGTTCTGATCCCACCGGCCGAAGCCGCTCGCCTGGCCAAAAGCCTGCCTAAC
TGCAAGGCTGTGGACATCGGCCCGGGTCTGAATCTGCTGCAAGAAGACAACCCGGACCTGATCGG
CAGCGAGATCGCGCGCTGGCTGTCTACTCTGGAGATTTCCGGTTAAATAACTTCGTATAGCATACA
TTATACGAAGTTATAGATC

SNAP

GCTAAAATTGGAAGTGGAGGACGGGAATTCATGGATAAGGACTGCGAAATGAAGCGCACCACCCT
GGATAGCCCTCTGGGCAAGCTGGAACTGTCTGGGTGCGAACAGGGCCTGCACGAGATCAAGCTGC
TGGGCAAAGGAACATCTGCCGCCGACGCCGTGGAAGTGCCTGCCCCAGCCGCCGTGCTGGGCGGA
CCAGAGCCACTGATGCAGGCCACCGCCTGGCTCAACGCCTACTTTCACCAGCCTGAGGCCATCGAG
GAGTTCCCTGTGCCAGCCCTGCACCACCCAGTGTTCCAGCAGGAGAGCTTTACCCGCCAGGTGCTG
TGGAAACTGCTGAAAGTGGTGAAGTTCGGAGAGGTCATCAGCTACCAGCAGCTGGCCGCCCTGGC
CGGCAATCCCGCCGCCACCGCCGCCGTGAAAACCGCCCTGAGCGGAAATCCCGTGCCCATTCTGAT
CCCCTGCCACCGGGTGGTGTCTAGCTCTGGCGCCGTGGGGGGCTACGAGGGCGGGCTCGCCGTGAA
AGAGTGGCTGCTGGCCCACGAGGGCCACAGACTGGGCAAGCCTGGATTAGGATAGATAACTTCGT
ATAGCATACATTATACGAAGTTAT

TurboID

GCTAAAATTGGAAGTGGAGGACGGGAATTCGCTAGCAAAGACAATACTGTGCCTCTGAAGCTGAT
CGCTCTCCTGGCTAATGGCGAGTTCCATAGTGGCGAACAGCTGGGAGAAACCCTGGGCATGTCCAG
GGCCGCTATCAACAAGCACATTCAGACTCTGCGCGACTGGGGCGTGGACGTGTTCACCGTGCCCGG
AAAGGGCTACTCTCTGCCCGAGCCTATCCCGCTGCTGAACGCTAAACAGATTCTGGGACAGCTGGA
CGGCGGGAGCGTGGCAGTCCTGCCTGTGGTCGACTCCACCAATCAGTACCTGCTGGATCGAATCGG
CGAGCTGAAGAGTGGGGATGCTTGCATTGCAGAATATCAGCAGGCAGGGAGAGGAAGCAGAGGG
AGGAAATGGTTCTCTCCTTTTGGAGCTAACCTGTACCTGAGTATGTTTTGGCGCCTGAAGCGGGGA
CCAGCAGCAATCGGCCTGGGCCCGGTCATCGGAATTGTCATGGCAGAAGCGCTGCGAAAGCTGGG
AGCAGACAAGGTGCGAGTCAAATGGCCCAATGACCTGTATCTGCAGGATAGAAAGCTGGCAGGCA
TCCTGGTGGAGCTGGCCGGAATAACAGGCGATGCTGCACAGATCGTCATTGGCGCCGGGATTAACG
TGGCTATGAGGCGCGTGGAGGAAAGCGTGGTCAATCAGGGCTGGATCACACTGCAGGAAGCAGGG
ATTAACCTGGACAGGAATACTCTGGCCGCTACGCTGATCCGAGAGCTGCGGGCAGCCCTGGAACTG
TTCGAGCAGGAAGGCCTGGCTCCATATCTGCCACGGTGGGAGAAGCTGGATAACTTCATCAATAGA
CCCGTGAAGCTGATCATTGGGGACAAAGAGATTTTCGGGATTAGCCGGGGGATTGATAAACAGGG
AGCCCTGCTGCTGGAACAGGACGGAGTTATCAAACCCTGGATGGGCGGAGAAATCAGTCTGCGGT
CTGCCGAAAAGCTGCAGTAGATAACTTCGTATAGCATACATTATACGAAGTTAT

List of parasites strains generated in RH-DiCre

RH-LoxP-CRMPB-3HA$^{\text{LoxP}}$
RH-Mic8extrac.$^{\text{floxed}}$
RH-CRMPB-3HA$^{\text{floxed}}$
RH-CRMPB-3HA$^{\text{floxed}}$ CRMPA-sYFP2
RH-CRMPA-3HA$^{\text{floxed}}$
RH-CRMPA-3HA$^{\text{floxed}}$ CRMPB-sYFP2
RH-SNAP-CRMPB-3HA$^{\text{floxed}}$ CRMPA-sYFP2
RH-Halo-CRMPA-3HA$^{\text{floxed}}$ CRMPB-sYFP2
RH-sYFP2-CRMPB-3HA$^{\text{floxed}}$ CRMPA-sYFP2
RH-CRMPB-3HA$^{\text{floxed}}$ CRMPA-sYFP2 Ron2-SNAP
RH-CRMPA-3HA$^{\text{floxed}}$ GCC2-sYFP2 Ron2-SNAP
RH- CRMPB 3HA$^{\text{floxed}}$ Mic15-Halo
RH- CRMPB 3HA$^{\text{floxed}}$ TSP1-Halo
RH- CRMPB 3HA$^{\text{floxed}}$ 247000-Halo
RH- CRMPB 3HA$^{\text{floxed}}$ 285500-Halo
RH- CRMPB -TurboID
RH- CRMPA -TurboID
RH -Mic15-3HA$^{\text{floxed}}$
RH -TSP1-3HA$^{\text{floxed}}$
RH -TSP1-3HA$^{\text{floxed}}$
RH-CRMPB-Halo CRMPA-3HA$^{\text{floxed}}$

Core sequences used for alignment of CRMPs

```
>CRMPB_TGGT1_292020 [Toxoplasma gondii GT1] EPR59691.1
CEKCAAGTYKELQENAPCAGRCMRDAETIEGAVAKIQCFCKIGKYAVVAEDAEGIITCQDCIAGGVCPGGLKTRA
RQAVEQDHSFVKITIDDHQVPFPSSGFYAVYKPLNETVWSPAMVPMVASFTGDTFKEFTDTGPAPDDHTDEGGAN
LDSTGGSGEPSSSAPVDPSGENEGQLLTGTTGAVSALQMRSRANDTSVHQYDRIPDIHPCVDDVRCRGGSHNSCV
HGSAGYLCSACEEDYSEIRYKSGCQACLPLWLDSFIFILMRLVVCGIIWIITALTIIAVQQQACIHPVLIRIVMS
HMFFLSVYGLMPATSQSQLAGWASIYRLFFFEFYFALHPYFKMPCFFRSLGIVMPEAHVWYWQHFFQIFVPFIDA
VLLTIIGAICVATYKVMYSAYISRVLVVLEEARQAHGDDMWTEKTIRKIESERCLGMFRYIYGTSTPWENFVRLC
TDLIPAYTAIWFWHFPTFVIECTMLMGCIETRYKSEDPISVLAAFPVQICSFENPYFLSGLILGGVGLLVWGVGS
IAGFVAYMSGDHSSDTIEQRFKHGFLVNGYQYAYRWWEGVIGLRKTCVALIITMYVHANASGAQEIFRNSANLAL
TVLSTALQLQLEPFDKRSHDMANRMEFYGLMVNIIIGVIFQGSYYFEVFKYMGAIPLAVAIFYYLYVLWSLFVEW
GRMVMMRPHLVSIPSLWRYYNRVTRSLARLYTSRNAKIYYNYITKDMVLEAATKSRVFHLRRLLLRRKKTSYRKI
NYENRTYFVAALSDSLSQLVIAWCQFTIPGDWLDFTQRYAFCY
```

```
>CRMPA_TGGT1_261080 [Toxoplasma gondii GT1] EPR62726.1
CEPCSAGSYKSSVQDGPCNGLCGTSSTSFPGAQTQSQCFCEEGTYFAADACHTCPIGAFCGGGLLEEAEAKLRED
SSFTGITSADHVKPFAEAGYFLNKLKEELESPNDWQFTECPIQNACLSHGVCSETMTEYLCSECRRGYTNTFSKG
EICTSCPSMVWNIVCLMGYYLATLLFNIVMTYMNVAAGFNRRSIHSIVIKIASNYLTGISVLSVIDFSTIAFPSW
ITDLTATVTETVSAKHSTRLMSVDCLLRDNFDLSFSESFFYTMVFYALIPIALPIVATIIMSIIVYRVRAWYHNS
TQRKLELLKQTMQYGLYSLAQQLKEKYEEDRVFMIFRYIALPGESIFRRAAKFMEDMIPIYVTVLFFVYSSTTRN
MLSLLDCTYIDFGRAHQAKYFLRAAMSVECTDILSGPYFKFFAVGITGLLVWSIGIPLSCFLVLYVNRKTLNSRE
TRLKYGFLHNGFVKKYWYWEMVVFARKFLVIVVSSVALIPSADKNGSRVWLAVVIAVIFLIIHLVTQPFDKRSYL
TLDKLENHSMTIWTITLIVLAMMIGSDFSGSVNMALLLFVAVLSCMFILEVGVSLMFAYFDNVRTQQTFFRVPVI
GYVFRFFARLSEKRRAREPIVVFDTENEVIQLVAAKRQTWTLFRALRKNINLAERNYFIKVMSESLGFAVVHMKL
DVIPGSFLEFALRLGLAF
```

```
>CRMPA_Cardiosporidium_KAF8821572 [Cardiosporidium cionae]
```

CRKCPRGYYKSTVDDVPSQCYCLPNFYFANQACLQCPVGAKCEGGFTEEATKALMANPALSTVSSSSHTKPYALK
EFYLHKLEEQLISPDDWKIIPCPIKYSCFRNNTCIESMTDYLCSECKLGNTNNFDKESLCKECPSFGANVTLLVL
LYIFQILFNIFMTYINVSAGYNRKSIHSIVIKIATNYLTCMAVLQVVRLTEIDLPSWVADATKNASKLGTLDFNF
FSTSIDCLLRSSSGSSHAESFFRSMLFYAMLPLLFPLTITFFVFVIVAFIKHSRKAQIAKKLFLLQQTKQFGLDS
VCEGLIEEYENDRIFMIMRYIPIPGESASKRIAKQFEDMIPVYVTVLFYVYTSTTSKMLSLLDCANINLGSVHGN
RYFLRAAMSIECKISPGSEYMKYFILGISGLLVWGIGIPLLSFLALFLNRRRLNTKEIRLKYGFLHNGYEHGSWY
WESVVFLRKFIVLIISNVDLFPTVSKVSNQIWIATVIAEIFLVLHLLIQPFDERSYSKLDTLENHSMSAWALTCI
LFGMVLNANLGGVFNLVLIVVVVSANTLFFGEVIYSLAFAYFDNVRKIRSFYDVPIIGVAFRFMGQLSSMQKEKE
ALVTYNSTTNDIELLIRRRKVTSKLMKFNQEKISSPERIYFKKMVQNLVALAVLTMKLDVIPGAFMEFALRLGLV

>CRMPB_Cardiosporium_KAF8819220 [Cardiosporidium cionae]
MCTPCLSGTYKRSHENTPCTGICMRSASSIEGAIYASQCFCDVGYYMVRDKKEPTLLSCQACIEGAVCPGGLRPP
VMYQILSHPTFSDIHFADHIPPFPKKGYFGSLRYVKKQLSKLLERLETAYVEVNYRERTSVGLRQRTLQPEMDRM
ALDLYEDRFISISPCPNPTQCTGGINEACKEGARGYLCGECQQGYDKGYFESSCIACSQRGVLFGRFFLARLVLY
LLTAMLVYASYNSQKTPMITHGALLRIWLNFIFFMTLYGTLPSTTASSLDIYIQLYQRLFAAPFLFLGSYFPFHC
VLAFTQTADEAYQRAWYSQRILQMSLPLLDCLFYSFFTWIFYLIFLLLHKKRIQQVTKAIAVAKEEASSYQAIER
VVSQMYSQVLWGMMHYLFPEGTSTKYRLRRFFQDLIPAYNIILFLHFPLLFVEFVQMSWCRSDTQNIAASDISML
LHLPSKLCNAKDSLFLLGVSLSSIGLLILCGLVAAWFFYELIIRTYQTVNRRLKLGFLLNGYRNQYCYYEAIQFT
RKALIAVTLAAHVQLSPRGPSEYFKITCSLFLTSAALIAQLLLQPLDSRSYNLLNTAETFGLIVNLITALIFQGA
YIYSFFKYVGLIPIALTITFHFWMLWMLLVEGGRFVIMRPSLVHLPVPWGWFNRLVRSLARMYIRGTAKVFYNYL
TQDIVVEASVQRTWFRFSSFYKPQAVYLSVKKSMREYFCSIIAESMLHLVLTLNRDSIPGDWLDFTMRYAFAY

>ParameciumXP_001426280 [Paramecium tetraurelia strain d4-2]
CLKCPAGSYSLDNFNHCQQCITSIESCPGGSELVLKQGYWRANKTTDSIEYCQSSVSQCLGGSEEFTCSQGYTGA
LCDDCDYYAEHWNNSYTRGFGGNCTLCQNDVFNIFKIILSFSWILIALFISIRGSLRLVRAQLSAHYLRMMGIFF
ATRSTMVIDQTEILIKLFSSYFQLISIVQVIDFDFPTPIEVIAQAIGNPLSTLGYSIECFLLNSQLQIEIVYLRQ
FWNLFVSLLFVFSFVFIYSLIQLCKQSQMENSIKTIFISCLIQVNFYFQGDIIEGLLQLMFCIKASGQYYILAAT
SYMCYTEEYYLYLKILIIPTLILVGIVSPLFYIIKLYRNKNKLWTCQLRMPYGYLYVEYKDNYYYWEFVCFFIKS
LFYLLETLLIQDIKLMFLFAILILLVYLELLNKHHPYIEKQYNFIDKISTQLAMVTLILSYSQDKNPYTSLVYIL
SILCSVLNLLYCTYIFSKIIREYLSSLKQQHIERIVNLIIRYPCIKYCIKKPKNYYNRRKACLLWKKVRVYVMEF
IEKLKLQK

>CRMPA_NCLIV_026200 [Neospora caninum Liverpool] XP_003882863.1
CEPCSAGSYKSSVQDGPCNGLCGTSSTSFPGAQSQSQCFCEEGTYFAADVCHTCPTGAFCGGGLLEEAEAKLRQD
SSFTGITSADHVKPFAEAGYFLNKLKEELESPNDWQFTPCPIQNACLSHGVCSETMTEYLCSECRRGYTNTFSKG
EICTSCPSMVWNIVCLAGYYLATLLFNIVMTYMNVAAGFNRRSIHSIVIKIASNYLTGISVLSVIDFNTIAFPTW
ITDLTTTVTETVSAKHSTRVMSVDCLLRDNFDLSFSESFFYTMVFYALVPIALPILVTIIMFIIVYRVCAWYRKS
TQRKLELLKQTMQYGLYSLAQQLKEKYEEDRVFMIFRYIVLPGESIFRRAAKFMEDMIPIYVTVLFFIYSSTTRN
MLSLLDCTYIDFGRAHQAKYFLRAAMSVECTNIWNGPYFKFFAVGIVGLVVWSIGIPLSCFLVLYFNRKRLNSRE
TRLKYGFLHNGFVKKYWYWEMVVFARKFLVIVVSSVALIPSADRNGSRIWLAVVIAVIFLIIHLVTQPFDKRSYL
TLDRLENHSMSIWTITLIVLAMMIGSDFSGNVNMALLLFLAVLTFMFILEVGVSLMFAYFDNVRSQQTFFRVPVV
GYIFRFFARLSEKRRAREPIVVFDTENEVIQLVAAKRQTWTLFRALRKNINLAERNYFIKVMSESLGFAVVHMKL
DVIPGSFLEFALRLGLAF

>CRMPB_NCLIV_012890 [Neospora caninum Liverpool] XP_003881528.1
CVKCASGTYKELQENAPCAGRCMRDAETIEGAVAKVQCFCKIGKYAIVAEDSEAIITCQDCIPGGVCPGGLKTLA
RQAVEEDHNFVQITIYDHQVPFPSNGFYAVYKPLNQTVWDPSMVPMVPSFTGDTFEEFTSGVSESASENDSETAG
GLDLVDESTEGSDETSGVVSPESAGGQGGELSEISGAVSTLQVRTRENTEAVHRYDRIPDIHPCVDDVRCRGGSH
NSCVEGSAGYLCSACEDGYSQIRYKSGCYACLALWLDSLLFILMRLVVCAIIWVITALSNIAVQQQACIHPVLIR
IAMSHMFFLSVYGLMPATSQSELSSWASIYRVFFFDFYFSTHPYSKVSCFFRHLGIVLSDTHQWYWQRFFQIFVP
FFDAVFLTVLGALCVAAYKVVYSGYIARVRVVLQEAREAHGDDIWTERTIRKIESERCLGMFRYISTTSSPFESF
VRLLTDLVPAYTALWLWHFPSFVVECTMLMGCVETRYKMEDPIKVLAAFPVQVCSLEDPYFLAGLSLGGVGLLVW
GVGSIAGFVAYMSRDHSSDTIDQRFKHGFLVNGYQYEYRWWEGLIGLRKTCVALIMTMYVHTNSSGAQEIFRNSA
NLALTVLSTALQLQLEPFDKRSHNMANRMEFYGLMVNIVIGIIIQGSYYFDAFKYMGGAPLAVATFYYLYVLWSL
FVEWGRMVMMRPYLVSIPSFWRYYNRVTRSLARLYTSGNAKIYYNYITKDMVLEAATKSRVFHLRRLLLRRKKTS

>CRMP4_PbANKA_1300800 [Plasmodium berghei ANKA] XP_034423161
CFPCPPNTFKESEYDNSCSGACPHFSTSFIGSSFENQCFCKNNYYLVTEKAHNIKNSNIKKSCELCPTGAICNRG
FNIDLYIKLLKDRAYNSISILDHENPYPVYGYYAVYKRNNKNTLWTPMDDKNSSSDDESAKNNTKKNTYTNFKYT
YPYYNLILYIQKHVQNMNYFFFENDNKFYLDKAFIESIKNKNEMKKNNNKHVDATKSNSDIYDQYNTTSFIQNAN

KYDTSKYKNDQSHNTRKEKNIYTKQNKNSNITHDPTKVDFYNLLSKQDQYLSFLESNKNVIKDDNNDTNNLIKEI
FNNNLKEAKKITERNHQFERTPDIHPCNLPDRCLGTITNLCYEGSTGYQCNSCSKNYDMKYFKSKCTKCRNIYHE
ILSIILLKIIYYVIIIYIVSLNYNSCLNNLYVSGVLFRIWLNCSFSFISLGFFSPNNLSFITRYWYVYKEIFLYH
LNFCSPYIRVGCFMSYYNTDITYKNIWYIQKYLNIFTFFFDVFFITLILFIIMKITNFWHRKKIQNFEIMLSTVP
EVYDKFKIEQNDKKKIKNKKTPKIHILNSNHEDQEDGSQNNLTLIHDLEKNESQSDEYATHGDAQNFQNNRNDCN
SQDEQGEEPHTLKNNTNVSVEKKINFLTKNKKSMLNNSQIENKLTEENENIHDENYELKYIKKEHTMYSQFLNEA
VEYIYDKKMFGPWRFIHKKNDCFRKRFLGFISDSIPCYILMIIISTPYILLETVQLFYCKSIKFKSEKSELYLAY
LNTQKCTTSSASFVMGLIVAFVVLLFYIATLILLLYLYSKRKTIKLFNKFLKNLLSGYRQGKEIFEVIFLLKNII
LVVMIAFTIYYQYYYIVLITLTLTLFSILELISDPFDRRSFNILNISLRVGSVLNIFFSIMIWGSFYLNYERHIL
FPFFIVILYHFYMVHNIVKEVILSKYFITIQTYINVQTMGYDDHTIKNKNYKFYYKVMNKIKFVLKKKHKESILQ
YAKNDSSIEKTHGQPKEYIHFEAIDNIIKKYNDNLKKTVIRNIDQNEEIIDNEQKNIKGKAQSSLKKKKLSVNVW
NVAYIWYDEQNEDLFFQMPFEDKLSSSFNKEPNEKTYNIFARIPDRIKDKETKYNIKYFVLAIIQVIDKFVVISK
SCSIYENWFDFAMRFAFVY

>CRMP2_PbANKA_0615900 [Plasmodium berghei ANKA] XP_034420659.1
CAPCESGFFKTIIGDFTCESKCKPNSTSFYGSTHETHCFCLENYYFKNGVCINCPDGAHCRGGFEEETLLNMKKN
ENYLDHSKIKHVMPVPRENYALYRLKANIHNYDWFIVECPIKDACLYNGKCYESMTNFLCGECKKGYTNNFSKLN
LCIKCSGIIANILYILFVNMFALLFIVIMAYLNVFTGTNRKSVHSIVIKIGINYFSCMKLFYIIGTSEMYFPINL
SSHVNYMTKYIKRLLKVKKNYGIYCILTSFNISHANAYFYGMLYYAFKPILLAMILTILMYIVVEMYKYKVRDKT
NIKLRVIDEIKVLGNNKLYDEIMQELISERGLVLFRYISIPGDSRFKRIKIFFEDMIPIHVTLLFFMHTEITYYM
LTLLNCKAIYYNDKFIEQYMSFAPSVKCDLSKDYAKFFILGISGLVVWGIGIPLMSYLVLYKNRRKLHNENVLLK
YGFLNNGFNFQFWYWETIVFLRKNMILLISTVSLLKTTRVLGTTMWLFTCVSSFFLILQIILQPFDSRNYHILNR
LETFSMVAWTISLIILAFLTVSSASSTVNFYVLLSLLFFNCIFMANILIVLCNSYMENLRHIKKNIKIPFLSKFF
EKMSKIAEEKCYKEPIVSLNLHDNSIEFTKKYKKHHLFGKRILTNEEKNYFLNVLSNFIYFGILNLNFTIFHSYF
MEFVLRLSIID

>CRMP1_PbANKA_0812400 [Plasmodium berghei ANKA] XP_034421095.1
CKPCPEGTFKNYDGDVKKCISCPPKSTSIKGSIYPNHCFCKNGFFYSKDTCLECLEGATCNGGLYPNAMKKIKLD
IENVNTIGPDDHVKPESKKGYYLDESIINIIDVSEWRFIKCPISGACLGKNNCHNTMDNYLCIECKKGYTNSFTK
SKCVRCPNNIANIILLLLIYIIFCFIIIIISYLNISSGFYRRSIHSIIIKIAVNYVSSMLIVNILEDTYLNLPPY
AYDVYNKMTNILTSENQKKKIISIDCLLRYYFNLTYNDSFFYTSLFFFLIPIFLMLTLTVILFVILKIYTIIQKE
GINNKLNLLGMAKKKNISFLVNSLEANYKKERFIMILRYIKLPDNTMSDSILTFFEDMIPIYATFLFIIHAKTSL
RMLQLFDCSYIRYTKKISKYILNSSSSVQCNFKTNDYLKFFILGISGTVLWVLGIPFLAFFVLYNNRHNLFHENI
RIKYGFLHNGYLPNRWYWEVVVFIRKIIVLFVTTVIVFPSNKEKIYKLLIITFVSIFSLCIHFIFQPFDKRNFFI
LNKLENFSLYIWVFTIMIISVLMHVNLNEFINLLVFFFIILLHTIFFIKLLICLFYECISNLRATFHFCKVPFIN
SCVKMLIQIAEAKKKQEPRVFYDKCSRQLAVVLPDKALMEGNRINIYAIVQSFFKKFIRSNYLQYSELNKFENQT
ISTYESEDKEKIKVVNLRSYENIENYGNCYNEMMFRNQKTDLSNFLLKNTNLSIYGKIKNEHRMFAIEIYKELLD
IFLKHVTFTYIPDTLFEFIFKISVNI

>CRMP3_PbANKA_0606700 [Plasmodium berghei ANKA] XP_034420568
CIKCKEKYYKDVVDDSLCAGLCMEYSTSFKGSISKKQCFCNTRKYMIRESSNEIKCINCPEGSLCIGGLKYKPLK
ILIKNINYTNIEVVDHTVPFPQKGYSATFEIVHPDFSWTPLNSVNLKINNYDNSDIVLSNKLLFDVFYTKRNYVE
LEVSKFINGNKTSLLSKNSIFFRKKIILNISNNKNNKEIYFSKQENKTPVLITENGEQIKYKKSFLTVDRIPDFH
ICPIMKKCVGGINNLCYEGSEGYLCSNCSKKYDTKYFRSQCFKCQKTKIEIFNFILFKAVFYALIFFLIYLNYFC
CIKKNFVFIGIFKIWYAFIISFLPYIFIMDSNSPREENYILYSQFFTSLPIKFITQHLKINCFINSYNNEKYIYV
WYVQRFIKIAEPIIDCFFLVCIFICFYIIYTWWNYQKISLIQKVIKKQINKEYNKEYYKYVSDFYYHYYQKGIID
TINNKRNKTYLEYMKYDCDYFSSYNSLSFRTPLLKKWKNKKCSKKLDKSKLHSVTSWKSNKNQSSKWINQKVSNK
SHKWINKKEKKQKNKTISLDDINFLESSSFSRNKELTKENSKEFDFLDVINFKKDISHKIKKTSSLPIEENKTKD
NYWTSICLLNIYNIKAMGIFRYIHPPNISIHKKIGSILSDLNAIYIIVFYIHFPFTLMSILELIWCQSTKYKNKL
PILILYHMPSQVCSFQNKLFLSGVLFSGFFFFIYLFLFIKYFYGTFKNFKVFQSYDRDFKCYFLFNGYNYQNRCW
DFVNIIKIVLFVTSFMCQFYTNKINNSKYFIFSCIIFIIITEITLILLYSPYDKRSNNVLQKLSLLSIFSVLITY
LTTHFSFFFDFYIISALPFILFIYFHIHTINKIVLEFALYKNILMKPTRQKQKLDSEEFLNENSILFNSYLNTKT
NIQDNDEDIIFSNSKKISRKYKRYKKEIHSKIKENYNLLNFVLRINKIPSCSIFFNEKNEEIFVQDSNDLIKCKY
YKNEKYFKKHNNSLKINPNIITLDILYSDLLYYNTTYSNNEIFTNLNNSPPPKYCYSSSGSKMSKDESLSNRNGF
REFSNTEKIVESSESYEEKNVLDNQIQKDTLKNYNSPLNKTHFIKCLTDAINILFVNQCYNHISVEWLSFVIRFS
ICF

>CRMP4_PF3D7_0718300 [Plasmodium falciparum 3D7] XP_001348896.2
CFPCPPNTFKENEYDNSCSGACPPFSTTFVGSSYENQCFCKNNYYLVSTDDENDTNGNSINNNKNIYNNNNIYNN
NNNNNNNNNNNFNNKSDVKKKSCKICPIGAICNRGFNIYIYLKLLNDRSYNNINVIDHETPFPKYGYYAVYREKT

NPTKSWDTLDDSAILNYVYPYHKLLLFIQKNIQKGNYFFLEKGNKFFLNDDFVEKMNEKSKYEEENANKFVTKSE
IIHNVNTNNLKGKHNTLNNNNNNNNINKSSSFLNQNNYNNDIIQLSSYENNPFKNLSSTSEVSSNNNVISFIQNN
DKTNNSTNDNNIENLVDIFNKYVQEAKELQIQNRKFERIPDIHPCTIPNRCLGTLINLCSDGSTGYQCNNCSKNY
DMSYFKSKCKKCQNLFYEILHFILLKLLYYIIVIFICSLNGNILNKLHVSGILMRIWLNCSFTFIAYGFFIPNMN
SFISKYWYLYKTAFSYHMKIFAYYLRVQCLLNSLNINLSYNQMWYIQKYMNICTPIFDSIFITILLFAVIHIYKF
FFYKKNIDIFEQLLCTIPELNTLYKKNCIDNIHKVKNKKFANKVESFEENENQNKEKKNKKNIKRKNKAKNKHNN
DNNDNNDDNDNNNDNNNDNNKININSKDEHINHFEKCSSITTILNNLELNNEEKNKESKKKKIFDIKKKTSIENI
KENETNHLWNDTHNNVKNKESNINTTNISNNINYELKYIKNKEWYNIFLNKTIQYIYKEKKIFGLWHFIFTKNDN
WFKNMGTFLNDTLPCYVLIMILSIPYILLELVQLTYCTPIKYKTQKTYFYLTYLITQKCSLSSKLFITGLIVTIT
MFALYIFLWILLLYLYSKRKRLNLFDLLTTNLLSGFRKGKGIFELIFLLKHILIVLLICFNIYYGLYYIVFLTLI
LITFAILEILSNPYDKRSFNILNKSLNIGNVLNIFFPLIIWGSFYWTYENFFIFPIFVIYAYHIYMFYHIVKEFT
LSIHFIKTENIVNIQTIGYDPENIKNKNYKIYYKVMNKIKYFLNKNQKKNISNINENKDKHLNKNNNSKDHVKFE
PIENIIKRYNLKLNKGILKDANNNNNNNNIKNMSDQVEDNKKDILFIKKNVLPVNVHNVASIWYDEQNEDLLFQM
PYQNKFCLNMNNEANENNDQINDLTKDKTSEEENNNDHVNDPTKDQTNDEKKKNKNKKNKNNNNNENNCHSNDNS
NNNNNKEVHKNDDNINHNNINNHNKSSNISTTNKKNYFNKFSDYIIDKRNKKSIKYFITSLMEVIEKFLDSSTHG
FIYENWFDFTIRFAFVY

>CRMP1_PF3D7_0911300 [Plasmodium falciparum 3D7] XP_001351985.2
CEPCKTGTFKNYVGDDTFCLHCPPKSTSLEGSIYPTHCFCKGGFFYSKGYCLECLEGALCKGGLNKNVMQRMKFN
MENILITEKDHIKPHSNKNYYLDKSVIDVVDVSEWKFIKCPIKGACLEKNKCHKSMNNYLCIECAKGYTNSFMTS
LCVKCPNNITNMILLFLIYIIFCFIIIIISYLNITSGFYRRSIHSIVIKIGVNYISSMLFIKVLEDSNVSVPISF
IHFYNKMEKFLHMGENKKIVSVDCLLRYYFNLNYSNSFFYSSVLFFFIPIFLIITLTIILFIILKIYQCVQKKQI
ANKLDLLRIAEIENIHFLASSLKKSYSRQRIIMILRYIHLDNYKTMDKIWIFFEDMIPVYVTFLFIIHAKTSLRM
FQLFDCSYIKYTKYLGKYILRRASSVQCSFKEDYSKFFFLGLTGTIIWSLGIPLFSFYILYINRHNLFNENVRIR
YGFLHNGYVPNKWYWEIIVFSRKICVLFIMTVIIFPSDESNASQLLLVTFVAIFSLCLHFIFQPFDKRNYFTLNK
LENLSLYIWVSTIICTSILLSMNLSKLINFIILFTLFILNLLFFIKLLFCLFFECLDNIRRSKACYHLPVLGTCA
KLLGDIMEKKRKREPIIFYDKLCKQICIMLPVQCGKVPKNKKNNKIKINHNKYIKNITYPFTRLFKNLSNSYKHN
NIKKKNNYMPLQENINENKNDINNMCQNQNTQSYVTCSSDLDKNNIIQQKKSHNDNNINLITENIKNETKNNVCF
YLSNSHNIPLYGGIKKEYRMFATEIYKELFDIFLKNISLTCVSDTFFEFLFKITINI

>CRMP3_PF3D7_1208200 [Plasmodium falciparum 3D7] XP_034009380.1
CVMCKEGYYKDVIDNSVCAGLCIEFSTSFKGSISKKQCFCTVGKYMIQDEKDENKCINCSKGSICIGGFKFKSLR
ELIKNQYYINLNIYDHSIPLPKKGYFATFQINHEDFKWKPLNSLHIEINNYEKGNIIITDKIILNVFHKKLHLIN
IGEKNKYDQADQGIYIKKNLNNNSFIHNNNNIKYIVEESKHLRDRTMANIPNEPILVNEKLEPIIYKNNFLMIDR
IPDFHLCPISNRCMGTSSNLCFKGSEGYLCNNCSNSYDVIHFRSQCIKCRKKKTEMINLLIHKFLFYIIIFFIMY
LNYYCYIKKNYLFISILKIWYLFIICFIPYSFIMEPIENHLKNYQSYFQLYIILPMRFIAHYFKLNCFFNNSYNE
INNNITDNKVHIYIYIWYIQRFLKIAEPIFDCILFTLIFFIIYVLYISWNNEKIQIAQSVIKKQIENIQNNSYGK
HIYDFYYYYYQNNVINIIKNTNNNCNVKNWSQSYYVHDYLSKTKKKKQKKKLSKWSLTQTKKKLIQRWKHKISRS
FKHKKENSENYDYFENKENKENNSLCSESTKHDNTINNNYTTSSVEDKFIYTKKKTTYNNNTNQSCDDENIYNVK
EKLLKKDITNNFGYEHIEKEKYWTSICLSNIYNIRALGLFRYIHPYNINIWDKMKRILSDLSCVYIIILYIYFPF
IIINLLELVWCQPIQYKENPSLLILYHMPSQVCDLQNKLFLSGVLYSCFFLIIYIILFSIYIYDTSKSIKAIGSY
SNNLKSYFLFNGHNYQNRCWEIINIIKVSFIIIPFICQLYTKRNVNSKYFICCCIVLVLITEVACILFYSPYDKR
SDNLLQKLSLTSSFFILISYLTVQVGFFFDVTILNIIPIFLFFYFHVYVIKQIILDFAIYKNILKRKDNHISFDK
KKKSNDMLNDYNFLFFFDKIEIDNKDSNTPKGIDTTVQCDEKEIKNKDDINNFNHNKINDVNNKRKKKKKKKKK
KKDETTKSPYNEEYNLLNFVLSSNNIPSYSIYFDDKEEEIYLFQNNNTRGYEYKKKIINNMKYNNDIKIEQTKRI
SLDLLHSDILYYNIRKGTINSDISNMNTYNNIKMRKQENNYYHQNDCNNINSDDYNNVCVQKRISNKLEHFIRIE
REYDKEKHYIKNNEKNNSEDDMFFLNITHFINCFIDAINILYINQSYDKITVEWLSFITRFSICF

>CRMP2_PF3D7_0718300 [Plasmodium falciparum 3D7] XP_001349085.1
CAPCERGFFKTIIGDFSCESKCKPNSTSFVGTIHETHCFCLENYYFKNGICLNCPDGAYCEGGFQKETLLYMKKN
EYYLDTSKIKHIMPVPKENYALYKLKTNIYNTDWFIVECPIKEACLYNEKCHESMTNFLCGECKKGYTNNFSKLN
LCIKCSGHIMNILHMIFVSIFILLFTVIMAYLNVFTGANRKSVHSIVIKIAVNYFSCMKIFYVMGISELYFPVTF
SSHVNYILKNIKRLLKAKKNYGLYCILTNYFDLSHADAYFYGMVYHAFRPVFLAIILTLLMFIVVEIYKYKVRNE
TNIKLNITDKIKELGKDKLHEEIMNELPSERALVLFRYIPIPGDSRFKRIKNFLEDMIPMYVTLLFFIHTKTTYY
MLTLLDCKALYYNDKFVEQYMSYVPSIKCDLSKSYSKFFILGLTGLIVWGIGIPLMSYLVLYKNRKQLHSENILF
KYGFLNNGFNFQFWYWESIVFLRKILVLLISTVPIFKNAKIFGTTMWLFTIISSIFLTLQVILQPFDSRNYHILN
KLETYSMVAWTMTLIIFVFLTISNTNVTINFYVLLFLLFFNFVFIAKILISLCYSYIENLRHMKKRIKLPFLRIF
FEKMSKIAEEKDYKDPVVSLNTHDNSIQFTRKYKKKYISCSFKNNMLTTEEKNYFLDVISNFIYFGVLNLNFSVF
HSYFMEFMLRLSIID

>CRMPA_TA16980 [Theileria annulata] XP_953733.1
CNACAPGTYKSSVMDSSCNGLCTQNATSFPGAESSNHCFCLMGYFYLEGGICSKCMEGAKCEGGLINYKMKTLKN
FVVNIDDHVKPVAVEGYFLDKINPKLRKPDDWKFIKCPINGSCLGNDKCSDTMEHYLCSECKKGYTNNFVKGALC
QKCPRMRPNISITILYYFGLLLINIVMARLNISSGYNRRSIHSVVIKIALNFGICTSVVNVVNFSDLKLPVGLRK
MINKWVKVFDSDDAVYYTSIDCVIRSLFNLEHSDSFFYTMLYMACFPLILLFIVTLMMWIIMVLFKLVKYNDIRR
KLALLQQFKAKYEGDNQDSLIEQYRNERLLMILRYIPLPNETSWDRFKNFLEDMIPIYVTVLFSVHGKITSRLLS
LLDCTYIDLGRSFPGKYVLRPAMSIRCSINPSEGYLRYLILGLCGLLVWGLGIPFLSFMALFVNRNNLYAPSIRM
KYGFLHNGFRQDYWYWETVVFARKCLVLVIGSIVIVPSENYSGSRIWMALVVAVTFLILQLIYKPFDERDYFVLG
RLENHSMITWTFTLILVSFIIEANFTPSLSLILFFIILFINFLFILELLFYFIIFISSHIFYLFSVFFPRFFSRN
IKISHPLSYKEPIVFYDKQDSHILLNRQIKPFGSLISFQNLSSVYSNYFTTTLSHIFNLTACYLNLDVIPSLFID
FLIRYSISI

>CRMPA_TA16985 [Theileria annulata] XP_953732.1
CNACAPGTYKSSVMDSSCNGLCTQNATSFPGAESSNHCFCLMGYFYLEGGICSKCMEGAKCEGGLINYKMKTLKN
FVVNIDDHVKPVAVEGYFLDKINPKLRKPDDWKFIKCPINGSCLGNDKCSDTMEHYLCSECKKGYTNNFVKGALC
QKCPRMRPNISITILYYFGLLLINIVMARLNISSGYNRRSIHSVVIKIALNFGICTSVVNVVNFSDLKLPVGLRK
MINKWVKVFDSDDAVYYTSIDCVIRSLFNLEHSDSFFYTMLYMACFPLILLFIVTLMMWIIMVLFKLVKYNDIRR
KLALLQQFKAKYEGDNQDSLIEQYRNERLLMILRYIPLPNETSWDRFKNFLEDMIPIYVTVLFSVHGKITSRLLS
LLDCTYIDLGRSFPGKYVLRPAMSIRCSINPSEGYLRYLILGLCGLLVWGLGIPFLSFMALFVNRNNLYAPSIRM
KYGFLHNGFRQDYWYWETVVFARKCLVLVIGSIVIVPSENYSGSRIWMALVVAVTFLILQLIYKPFDERDYFVLG
RLENHSMITWTFTLILVSFIIEANFTPSYNMSLLSFIVFINFLFLFELIFQLFVAYCHNFRIQNTSPKSFFTRFF
YHFLTKISNKIKNAEPIIYFDPSTCTTNLRSSKKFINKSIFDNIDVISYNEKDYFSYVMNEIICFANIRMKLDVI
PCDFPEYITRLTFAV

>CRMPB_TA20783 [Theileria annulata] XP_954318.1
FKEGCIPCEPGMFKEGQENSPCSGRCMEYATTYGGSVSKSQCFCQYGRYMRVKKQNNKVGDENHSFNSSENPAGD
YKNIKCSKCMEGATCPGGLNIEIVKILKKDYTFDSIKLSDHVKPKPKQGYFPAYPAYNNVEKKSKIWNPSEQNVS
IVERVYTDFVEFHPCNLEFRCGSQFQGYCSRGSDGYLCSRCMRNYDIWHFRTDCSKCKSILREIWSYLSLKILIY
SMVLILIVILYYDQIVCFVLIKIFMEFYFTMIPYGIISVFTNSSIKRFVPVFNSVFGYPLWSFAYFRLGCIFKSI
NYVSLWYFQRLTSVLIPVIDSIILFSICAAFLMLIERTYKITSSTSEMDTFGRDYMSEKKWRVLYFRILAICYFI
QLHFIWIELLQLIWCVNLSYKRQNHISVLLYMPVELCSIKNFRFVLFATISGFILVLSFCCFCYYISNHKLDRVG
EIFSFGYRRGFREWDSFVLLRRFIVSIICTIQIGIISPLESGRLRIISSLMLSCLIAVFNFTLYPFNLRSDNLFN
RMQMLGISVNITTACIIQVLVCEIILFRAVSVTTLTLAVWRIVDFFTLWHTNAEITLNLKKKCITVEPPFEVLKG
KFEKYRRNTASLSISSRNALSLSIRESLNRLILEKKRTFVPSWWLEFVVNYTMCRY

>CRMPB_TA20781and2 [Theileria annulata] XP_954316.1 and XP_954317.1
CQKCLPGTFKLVYENEPCHELCPVYSTSLPGSESEKECFCIPGFYNNFKNGKFQCKKCPYGSICYGGIQNGIHTK
PVPKSGFAIVDINNSRNESTNSFLQIFPQNTNPVVKNNHIRLTYPCVTPARCIGGGKCFEGSSGLLCTECDPGYD
LHHFNSKCKKCSSNIRELIKIILPRITLYCLIILLSLYNKRANSTSELSLVTIFKILYSYTLSLVPLGIIPSNSP
SSIRKFYNFYEKFFYHPLNFYAYVDRVNCFRNLAVYVDFILNRLGFDKRVPKLTELNYIEIWYLQRYFGIFKLIL
DIIFIFIIDFILYIVNKTFSICQRSLKRRLKFLRIKVGTSNVSLSTSHDETEIKFETDPKLIFQHIVVLICLHLP
SISLNSLSMLWCTKFPLRKGYVLLHMPNQECTFKNKYFMFGSIVSGSSMGFIFILLLFLFFKFWNCEYTSGWGNI
FITGYRKKYRNWDVVQLIRQILIIFLIVCRNSIDSNKSEINRILFYLIIHIIYLVIMLSTKPYDSRSGFVFLNLE
KYLIASNIFSSMFMYGSYFHNFSIISGLPFIVSAISYVLIFSTLLREFITICCIISRPREYFWRNFTSSFISFFK
HNHSLLYFNQHDDTMVFEAYNISKKKSIDSIIDSVNIKRDKTFLITCLREVIEVCISHKNIGVFSSTLFYFYIRL
IFWN

>CRMPA_BBOV_III011730 [Babesia bovis T2Bo] XP_001612293
CVPCEPGTYKSSVMDSSCNGLCTQNATSKAGAHSPAQCYCQRGYYYLAGGICAPCVEGARCDGGVVEVDEVGADA
DGNDDQDDDVVVRRTHINPVPIEGYFLDKINQELRKPDDWRFIMCPIKGACLGEKGCSESMTAYLCAECKTGYTN
NFKKGTLCTKCPKTMTNVVLTIAWYLGLLLVNIVMACLNVSAGYNRRSIHSVVIKIALNYGICMSVLNVINFGDL
ALPAELKSYTLRWFKVIHRESSVVHTSIDCLLQHWFGMTHAVSFFYSMLFIACLPVILLVVVTVLMWVILELFKI
KRHHVTRSKLALLYQSSLQGMHYLADRLRDEYANERLFLIFRYIPLPGETRWVRFKHFLEDMIPIYVTVLFSVHG
NTTSQMLSLLDCTCIHLGQSVPSKYMLRPAMSIRCSLDPGSAYFPYLLLGLGGLIFWGFGIPFFSYIVLLVNRKK
LYAPDVRMKYGFLHNGYQQEYWFWEAIVFTRKSLVLVIGSIVIVPSENTSASRMWMALAVAVVFLIIQLIYKPFD
ERDYFVLGRLESHSMISWTMTLIFSLFAIEGEFSAYNNMYLLSITSIINFCFISEVALELGLAYCDNIRFRKDQC
STSIISKINRSLAAISEKRKSREPLIVFNELNRTIDLCAPKSNQWYVFDERNRRLTYSEKVYFTHIVTEVINFSN
IHMKLDIIPTDFPEFITRLAFAV

>CRMPA_BBOV_III011740 [Babesia bovis T2Bo] XP_001612294.1
CVPCAPGSYKSTVIDSSCNGLCTQNATSKAGAHSPAQCYCQRGYYYLAGGICAPCVEGARCDGGVVEVDEVGADA
DGNDDQDDDVVVRRTHINPVPIEGYFLDKINQELRKPDDWRFIMCPIKGACLGEKGCSESMTAYLCAECKTGYTN
NFKKGTLCTKCPKTMTNVVLTIAWYLGLLLVNIVMACLNVSAGYNRRSIHSVVIKIALNYGICMSVLNVINFGDL
ALPAELKSYTLRWFKVIHRESSVVHTSIDCLLQHWFGMTHAVSFFYSMLFIACLPVILLVVVTVLMWVILELFKI
KRHHVTRSKLALLYQSSLQGMHYLADRLRDEYANERLFLIFRYIPLPGETRWVRFKHFLEDMIPIYVTVLFSVHG
NTTSQMLSLLDCTCIHLGQSVPSKYMLRPAMSIRCSLDPGSAYFPYLLLGLGGLIFWGFGIPFFSYIVLLVNRKK
LYAPDVRMKYGFLHNGYQQEYWFWEAIVFTRKSLVLVIGSIVIVPSENTSASRMWMALAVAVVFLIIQLIYKPFD
ERDYFVLGRLESHSMISWTANLLAVSLMCYVTLTPFVVTTVCALVLINSGYFFYVVFKSFLRSLVDTIRYKGENR
FFKKLRWLVDAVLFFDSRRYLREPQITYNQTTSYLQLRITRHRYTWFGKKTYELKPLERYYFVNIVTELYRIVVS
HLKLDVMPKLLIEFLVRLSLSL

>CRMPB_BBOV_IV006250 [Babesia bovis T2Bo] XP_001610552.1
CVKCEAGMYKEGQENSSCSGRCMQNATTYGGAVSQKQCFCNYGRYMTLDDTLGSMQLRCAPCLKGAVCPGGFFDS
VIEELKGDRTFTKIRLHDHRRPMARFGYFGAFKHQGIDLWSPVTQYPSSLNQISTDLLDFHACPLEYLCTSDNAM
PCDVGSTGYTCMNCIDGYERSYFRSNCLPCSGIPNAIYIYLKTKVFLWIICSAILYLLHKKLYYQFVIIKIWLEF
WLSMVPYGLFPLNSSSSLQRFSTKYNMIFGYQQRFFSYVRVSCMAKRFFNYNLSTFQSWYAQRLLIVVQPAFDGL
TLYFMLLILRILFYPIGKVRRRLSRRYSTWSSSGSSVSSYDDSASWNATTKTNTNETNEIVSGEIVVTNKKPPRK
RSLWLGAPRCILVMYYLSFQFICQELLQTLWCVPVQYKSEPPISVLLYHPSTVCDFKDRMFLFSIITSCFLLGSL
FLTSMFLVFLSVRKRGYMHLFSSGRQRHTLSWDAVLFFRRLCVAVVVIFQPYTLSTGSSEKLRMIGSLLITTVLL
ILHLSVLPYQVRDDNLFNRLELFSMLLNSSTGFLILGSFSYDFNYTGLFPLGASILYSCLLLWYMLVECGFIADL
RPALRRRFGIIPNFWSFCRHLLLWHCHSHITFNYKSAEVVIERPFTGHTSSINRAVNNKRHYHVSTTARRGMCLC
IEQSVNRYVMEQRKFCVPIYWEEFLVRYSFAY

>CRMPB_BBOV_IV006260 [Babesia bovis T2Bo] XP_001610553.1
CQKCAPGTFSETYSTSTCHGRCNDYAISFPGASSFKDCFCLPGRFMTLQLIENKEKFVCKKCSDGTICPGGFTVR
QSGATIPDNLKEAIFSHTPQYPRMGYMAIFKRQRAEGSDIQWLPNKHDTYINEPPTPSQFGKYEHVADIHPCVYS
NRCNSFGKGGCSEGSYGYLCGKCAPGFDARYLQSDCVRCKHPLFELLDLFVPRLILMFLVCLICHMNNRATTDGN
FSIIAIFKIWYIFELSLIPLGMQQLTTSSSLVRFYNLYHNYFFKPILLYVHNERLNCWRPAIHRIWQILRSWGVP
LPPADTHEDLDYNHIWYMQRFLGLAKPLIDALLLCILYYPCRLLYSRVESWSASSSRRISLDERIFSLTRRVNNE
NEEIKDKEEAALCLKTLDDPSAHETIGMAYIKHRKRGKVFLIQQLMLLMAFHLPSVVVNSLAMLWCGSVKYKDGD
AVKVLMHLPEQICDPNNRLFFIGRIVGLANLGLWLLMLAVLFVALANYKYGSRSWNTIFMAGSDTNCRWWDAVHL
SRQFLIAVLIVCHYGIRPKGDTEYMRAACYFILHFIYVTLHLTFSPYDKRNNESFNNLESLVLFSNLCMCIFIQG
SYFYDFSDFGGFPIVVSLLVHVKIIWTIMLEYGHIQFANMKNNGKGELANMIIDFFSFAFEHRTANVYYDYKNDN
LVMESSKIQEPYRSFYYNLVTRPLLKLAHLTGISMKGTEKRKSYSYSLQNRDFLVECIRNTIQRCSVLRQDVVLS
DKWFYFITRYVFWY

>DICDI_Q86I19 [Dictyostelium discoideum AX4] XP_643780.1
CTECRTGSYGYNGETCVQCPSIHAYCPGGAKVNQFEGWWYDVNQTDKPYVLYECPNVNCIGSNTSDMCAPYSFGP
LCAQCDPGYYDWGEGCKRCDKTSPILFIAIVAVSIGMIVLVQSSDSESGLMTVIIYFAQTLMVLSTGVKFNFLAL
LNLEFQPGGSSSIFGICPGPFDYYQQHYFTLLSAPWLIVVLLISTVFFYIVYRYKLIQKLKNKIIKEQEDIEKSK
SISNYSSIIVTSDPGTINNSSLQKSLERKSLLRKQQQQQQPLLENDDDDSFDSTNTTTNDGDLSRDGYSTIADGG
KHSILGKSIETSISGSLSSQVTLIDNATAIPSNQTFLSQQLGAFIKLMLNIYTPIGKATFELFFCENVGVSAYVL
IANNGVSCYTDQYHKALMVSYCLLIVIIGFPLILMILLFLNRKSLDDPHTQRTYGVFILKYKSSFYFWDVILLLR
RLLIILMSTMDPTSAARSFLLVVVSLVSVLLQLKYQPFKRISDNRLELTSLSLLFICCVYLDNTVYQSIEQWIVI
SSSIIFFLHAVYILYSNYRYQIIDSLQHYLYIISRGKKGRKNSNTTLFGSNSYFLYNQDHIYNDDDFSRDSGIFS
KKDKKQSRTDKIVKKSKINNNNENNIENNNENNENNENNNNNV

>CRMPB_BESB_019000 [Besnoitia besnoiti] XP_029215968.1
CAKCASGTYKELQENAPCSGRCMRDAETIEGAVAKIQCFCKIGKYAVQAKDAEGIITCHDCIAGGVCPGGLKTRA
RRAVEEDHNFVDITIDDHQVPFPSTGFFAVYKPLNATTWKPSMVPMVASFTGDTFVDFEAPPPEDEDLESLPGSL
SGAAASGEPGVVIPPGSGQLPSGEPRRSQPKEDAVVSALQLRVRDVDFIVESARYDRIPDIHPCFDDMRCRGGPR
NTCVDGSAGYLCSACAEGHSQVRYKSGCNACLSVWLDTLVFVLSRIVVCAIIWIIASLSIVAVEQQACTHPVLVR
ILISNMFFFSIYGLMPETTQSQLSEWTGVYRVFFLDLFFATHPYLKVSCFLPSMGIELSDAHQWYWQHFFQIFVP
FLDVVLLTILGAIAVTFYKVVYSRYIDRVRVVLHQAREAHGDDTWTERTIRKVEAERCLGMFRYISADDMSGWRT
FLRLLEDLIPAYTVIWFWHFPTFVVECTTLMGCIDTQYKSEPPFKVLAGLPIQACNIEDPYYLAGLCLGGAGLFV
WGVASIVAFIAYMSTDHSSDTIGQRFKHGFLANGYHYEYRWWEGIISLRKTCVALIITMHVHLNSSGAQEIFRNS
ANLAVTMFFIMVQLQVEPFDKRSHNLANRVEFYGLLCNIANGIIIQGSYYFELFKYSGAFPLAITAFYYLYVIWT
LFVEWGRMVMMRPYLVNVPSFWRYYNRVTRSLARLYTSGNAKIYYNYITKDMVLEAATKSRVFHLRRLLLRKKKT
TYRKINYENRTYFVAALRDSLSQLVIAWCQFSIPGD

>CRMPA_BESB_084890 [Besnoitia besnoiti] XP_029217299.1
CEPCSPGSYKTSVQDGPCNGLCGTASTSFPGAQTQSQCFCEEGTYFAADACHTCPKGAYCAGGLLPEAEAKLRED
SSFTGITSADHVKPFAEPGYFLNKLKAELESPNDWQFTRCPVRNACLSNGVCSETMTEYLCSECRRGYTNTFSKG
EICTVCPSMVWNILCLTGYYLATLLFNIVMTYMNVAAGFNRRSIHSIVIKIASNYLTCLSVLSVIDFNTIAFPSW
VTDLTTTVTESVSAQSRTRLMSVDCLLRENLNLSFSDSFFYTMVFYALIPIVLPIIVTIMMSIVVSRVRVWYRKS
TQKKLDLLKQTQQYGLYTLAEQLKEKYEEDRVFMIFRYIPLPGESIFRRMTKFMEDMIPIYVTVLFFVYSSTTRH
MLSLLDCTYIDFGRAHKAKYFLRAAMSVECTEAFSWPYFKFFALGIGGLFVWSIGIPLSCFIVLYINRRTLNSRE
TRLKYGFLHNGFVKKYWYWEMVVFARKFLVIVVSSIALIRSEDMNGSRIWLASVIAIIFLIFHFVTQPFDKRSYL
TLDRLENHSMGIWTLTLIVLGMMVGSGFSGNVNVALLLFMAILTCLFILEVGISLMFAYFDNVRTQQTFFTVPVL
GYVFRFFARLSEKRKAREPIVIYDTESEVIQLVAAKRQSWNVFRRRRKNINLAERNYFIKVMAETLGFAVVHMKL
DVIPGSFLEFALRLGLAF

>Cvel_14651 [Chromera velia CCMP2878]
CTECPAGAFANSTGSSSCTECYPGSYSADTGNVRCSKCQMDFYAPVGGSESCLTCASGAICDSTGTVLPATAANY
FRYGPDTDPLYLKCPVKGACLGEEENECYKGSTGFLCGSCEENYERWIAPLKCQACPATTIAALVIVAGFAAYSL
MIWYLSKEAESAGLDKNDITSVVLKSILNYVVLLAVVFVAAPTSYPKWISEIVHAVELYERPFAWLSFGCLLRKE
FAPTHEILSKAVVAFIFPVVAVFLIFLAQFLYSIIAPLACKRNMRNHEISCALPTDLAVFAEKRATVAAAAARAQ
ALNGRGEGEGDDDVAMSVYGQQSRMQSFAASERSRSFSALLADPLFEQQGEVSESRGRPGGMGGFQGLSDAQYAK
WRRNTVEDLSPSTRQQLRSHTLQEIRRQHSEKQQQEEAEEELGGEGEGEGGRANAVAIADEGASINISRIFSPKP
RNEKAPEKEKPPETEEEQVERVLAEIEATDDADQHSASSAPLLLEAGRFRWNAAKRILRGAVPTLIILFFVVGGF
FWGSFHAILFFVVGGFFWGSFHAILFFVVGGFFWGSFHAILFFVVGGFFWGSFHAILFFVVHPIINVVLADTLAC
KYYEEVDRTVHHNDSELDCRGETYRRFLPLVYSGIAIWGFGIPLAAYAILFFNRHHLQSTWCRLQFGFLYNGYQL
RFYYWECVIMAREVLVHYLLVLDIEYESKLTLFVVVGLILVTLQAAANPFDSRAYGALNTLEMLSLWCFTISAIG
NLFWVNPTGNQNVDIALMVILVGVSNLLYLLYAAGLLFGGAKFQRAALGFSRMISPMTWLGMCISRKQREIKVPR
SNSRGLLRKHSFFDRFTQRARRASDALLRRGSKVGQEVSQRGEGENQNPVSPASSVVSPGDEREQMRKRGRTRST
VQIKIEPEDLEAQKGGGNLEGVDLEEGRLVEENAEGACPV

>Cvel_10473 [Chromera velia CCMP2878]
CAACTSGTYKEAEANSLCTGICQDNAETIAGAISEIQCFCRADFYFVQGTEDSSRFYRGTECRACPTGADCAGGL
KTSISSAISSNASFVNVALSDHNLPVPQAGYFLTTNNSLDLAEGKSEAVIAKCWLAEACLGGESNVCYGNQEGLL
CDECKYGYTRVSTANVCASCPNQAVNVVIIIVAVCVSLLVIVVLCKTTLQSGLDRKKLHSIVLKIGTNYFGLLSV
FQDYNLAELAWPEWVYNVFNFFLSSSVGDPFSFVAVDCLLRKPADLFVDVLFLMKVLIFLAPIVIPLFVTGLAAL
FVALYRNRHRGELQRMREMLDALEDSGPVFAALQHNMRTEFRDRRFLGIWRFFYEDDDDWKKQVSRYVERFWYWE
AVISIRGVLIATSSSFYVGKDDAIKLSFTLLIAVVALGLQLVFQPFDNRNGSVLNKLENRCLIVWTLTVVVFQSI
YAFNVGANFNAFMAFVILALNLYFVGFFCLIFYREVCNTVKANPELLLLPQPFKTVTWAMARSADNFIDNQGKLF
FDSDTTDVSLRKKSEREKERERNGGGKEGKTKKDKARERAGSVISLKSRGSFVGSELKSEGGRSEDENDFDRKKS
IVTLMADKDREAPRESRRMTTLGLNEKELARHKKVETRQLLGLSFGERLYFVAALAEVLSHVVVTLRMDSLPGDY
LEFVLRLAFTA

>Cvel_18313 [Chromera velia CCMP2878]
CRTCPPGTFADGFASDSCKPCLESALCGPTEAERLEAQRNITSMEIASWGTQSDVLQERLAGFNASATDAYKYGA
WAGLAELRESQYKHLRSICPSPREGFHAVAVRDNEGVSSSAAERRGPVSLCACVYDWLRYDGLGRDEDFFEAALK
DIEEQKDREEIGERGEIRGCNRFFQSLKKSRSQTHAPIVEGTCPAAQVCEGYSQNSLCHLALEESTSPLQVGQEE
MGFLCGAEDRNPRDWGDSTDLSLSDLLAGRGLQVQCQHRDFLHSQSLSTNMQVTQNMMGSGEPSAQATAAISRAE
SGGGVSGAFFWRCPWGDTVCGVDNECLLKTTGPYCSRCEEGYTRSALSSGCQSCDKERGWLIAKLILILIFLVLV
TSRVVKCILTGRGRIKTAERSMQAAVFKILTTWFELFIVSMKLMLDSVAHYRSLLEKNGPFLHGFVSSISLSGGV
PIHQVFDSPVTFTFGLDCIPELKTALGTDRPEQIHLMVALMFPLMALAIAFSVAVLAQLCCRRDSERRRAGLNRL
LLSREGAMSAGSLSGKESRGRGATRDGEGGRRTVSTDSSLSADGRGSERGRSRSPVPPPMKSESEDPIDFDQTSA
DLDIETGGRGWNDQRANTSHSLPGGGSVTLSRAWTGPIAKPAQKANDAVPNFRRSETVGEATVMSQNSDLLCNSI
QSGRSGKNLSFWRTLFRSKAKRGGDSVTGSNQPSEAASSVSGTEGRTQKASPWARWRSRVMNRCLSSWVVVLFLS
HPCLAREFSRNVSCVHVNRGNTFDNGDGKGRPLFLAMDPNLSCHHGPHQKWYILAWIGLFVWTIGTLTGFAIILR
SIRFRLHEDRVLTRFGFLYRHYKPRLFAWELWAMSRKILVLLPLWIPMVSSLRTTLWLVIAVVFLWVHALAQPYS
TSCFSICNRLEQHALMAFFVTILGAQVIGQLGVGVEDLLEQGRNLPGIIVSLNCIAIFVIAVLSVHVGFFLRVIL
VLCRPILRQLASASRLRKKFHCGKWMTSGWKRYRKSRRSGWARPLIGFGVEAETNNAESENGDRKHAQEPVKKTR
SLHARARLLWGFRWSTDDVDAPAQCSCLNRRGPRRVSMSSWFGACMEKKLRVVRVTLYLLLLAVEVLFESSCSLC
GFCLCWSRKGWRLWGWARSLLLLQQNPVVLREGRKV

>Cvel_7366 [Chromera velia CCMP2878]
CTPCGSGLYKEAAGDTPCSGRCMTNGQTAGGAISLRQCSCMQNYYLKSGPDYTEFWVGASCEACPTGAVCAGGFS

NDTLAKLEADGDYVEISAADHMLPEPIEGYYRLGENATDMVFDINNAAIWPCYVQRACLAGADNLCHSNNDGPLC
DECMPGFTRVSANNVCAVCPAGIWNLLLVLSIIVISFFIVVFYAQLTLQAGTNKKAMHAIVLKIGVNYVALLSTL
STFDYSTLDWPDWTTGLLSSFTDASAGDPFSLPAFDCLFRSEQNPVPFTQAFHYTKLFIFLMPIAWPLTLTALMA
LVIVVYRRMVRVRLEETRQMLERLRELGPLYGPLHRRLLEEKREERFLGIWAVFSLRGERLIVQARRFMSDTIPV
YIVTLFLMQNSVTTHMVQLIQCDYINKERRLRYAVSQTCDVTDPFYVIGWMGLLLWSIGIPVVACSLLWKHRKEL
WKPYVKRRWFCAAQSSVGSGSVSLELACRYGFLTNGYDEGFWFWESVVTVRRVLIVIAASFFVGTRTNIRLLFMI
IVATVALALQLTCQPFDNRNGQILNRLEDRALYSWTFTLILFAMIFIFNVGSWFNTLLVIIIVILQLSFLGLFAH
HFYREMCNTVLNNPAWWDWPIPLRWVTRALKRSAEKFDARQARLTFLTESADVLLGEARTGTSSWVRQRRQTEAN
VFAARMKSPTREGAEMNESAKAGRNRESVASARQASFHRKYLSASGEGLGVQEEEPTSAAPGEEGKTEKGEEEES
DANTSGGFAHRSHKVHTRQCTISLLQDSDEDDAPSGANKLKGRRASSVWESNPFLKFVRRRICCCCKRNGTSGKD
EDAPPPLLKSDRRYFAHMVGEALTYALVALRLERLPGDYIEFILRFAFAS

>Cvel_29734  [Chromera velia CCMP2878]
CELCPEGLYKDLVLNGPCRGRCLTNGATVPGAQTARQCYCKEGHFFLPGPDANTFWEKAQCKPCMNGAYCPGSLR
APILARLKSDTNFTAISVDDHGLPKPLTGYYRISADPEDIHQCPSPESCEGGAVNSCSEGHTGTLCGDCLPGYFR
LLGQTKCQRCAENRYGPMVSIGFNAVVCMVVCFLFGAMVVWCSREVRDGLSPLIFRIILNFIGTVAVLQHFNLGQ
ILQSKMQSWSASFISILLNLDELNPFLNFNFECVLRAEALQLGAAREIRASEDAYLPASDVFVFTQIFYLVLPFL
LLLGLVGISACLLGLLHLCNRRHVREQSLIYRGLCDIVAGSGPSPFDTTDKKLQMSGKEGGAGGPSVSNNLNATD
VQKAQADSMKLEAFRRTVRRRFKVQRAFGLFRFVYKTGSFVEILGLFFEDVAPLCVAALILLHNTVSAQLLKLLD
CVQLANVQTLTDSASATAATATTGTQVEDTDGFAYELRMRYALEISCSSDRYKVFRNIATMGLVMWTIGIPFFVF
VLTYIKGRKHMAGKTKTTEGSSKSKANLKKKMNSKKANTVLKQQEAKEAMRREQAKQQADRFVWSFLWNGYEERY
WWWELMIFARRACLLLSVVFFPASVDASIRLIFVLMSSMVFLAGQTMASPFDKRSGGVLNGLEFLSQSTALAVFF
VFQLVYVWPVDNSFIEGGMLVIGILCLHFYVRAALALWGATLEQARLADPSKVLTLPFPLNWVLVQLCQMGNARV
RSQPRVWLDTQSRSLLLLRSMHKAANSVVEFLHTLAVRKGGRLSAAKKNRLDEATLLAQEAGDNAPLSDLGPDDE
TRASLLLKRSRRLSEQERASAYTLEDEEEEEEATMGEEGETVEDDERSERTEQSKTTDKEEARRANKKREQESLR
RSKSKSLTKRKRRGDDDGLTDALYIPPFNGKARGFLVTVLTEALEYSLIDLQSPLVAADWLEFLARIVFAD

>Cvel_1447 [Chromera velia
CCMP2878]CEKCPRGRYKEEVGNSACKSPCMDNGDTEPGAQSLRECFCGPGTYFEPGGNKEFFWLEASCQPCVD
GADCPGGFGDEIMASIRAKIADAEDKTGPLQSTLKSESSTTTGGTATGTTGRRLLEQEDLFLPAEEEGEGTVRRL
QAEDVTLAAHAPPNPLPGYYRVGTTSTILPCKVADACTGGDASDCTESMEGLLCETCRAGYTRVSIEAKCAKCPE
ERWNMMANVALIIVSWFQLIIFGRMEIESGFTRKTLHSLVLRIATNFIDATSVFRSFNLDRIKLPDTARDVFSNW
FQGGSADPTALVSLECVLRPEGSDLSQANALFYNQLFQFLNPMLSPIIITLIAFMAVAIQSRLHRQKTLSEQRIL
TDLAERGARYGALYFLLRKEFCGRRVLGIWRYFFEAGTSLKARVAAFLADTLPFYLVCWFQMHANVTKQMLKLSA
VKAECSHKIQCEDINGETRLRHATEVICNSDEHQPWFILGFMGLSLWSFGIPIYAVFVLFKQKKNFHDDFVRQRY
GLLNNGYDMRYWFWYSASVIRTVAIAFSTVFYFGEDPSIRLGLLSMVGAIAMILQLKMRPFDKRNGDVLNRLENR
FLVSWTLSLLLFEMIYLYDMGSVVNTVIIVVLLMLNVYTMLLILVNFLREFSNNIAGNPGLVNMPGVLGALVRAI
IAPTEKSKAREPKVFFNTVSGDLVLDHPVRVRPNTPGGRKSVERDREGAETPAGTNPFGFKRSQTANFSIYDESP
VSRRGGASPKSPHTPGSVAGTPSMAGGGSKAVALKDRRPNSKKDFFSFQTNDRRYLVTALGEVLAFLMQSLQLDT
LPGDFIEFLVRWSFYT

>GNI_010550 [Gregarina niphandrodes] XP_011128770.1
MSCPTGYAGPECAYCDEGYWVDRSALKGEQCHRCPGSVALYMALALVVSTLFCLSLAVLNVKAGANRKSIHSLVL
KLMIAFVADLQTYTTYLTTTLLHYVNLGGGSDSSSTPSGVRIQDLYDLGFLQCLSKSDTFAERYRDGMLFFNAVP
LLSGLFVTFIAASVVGLSIPLHANSIRKRRRLYEMLIQKNLIDAAAKLGREIDNTRTLSVWRYTTARPIPVAAHV
IRFFRDMVPPYIVIGVTYYNILLRRQTDLLACTRDAEGVLRNASVTGIKCAWDDSNYVVYFSFGLVFMFFWGLGL
PVVFLCLLLPHSNQLNSPQILQKYGFLHNGYDLRYWYWEIFFFARKAAINISLIISHNDSNYMKFALFISLMFFT
IHVWISPFDRRNNNIMSWLDIGCLACWIVSLTLACFGEALDTEGSNVIGLVIIFINVIFARTPAHSRTVPTYATY
T

>CPATCC_000266 [Cryptosporidium parvum Iowa II] XP_627992.1
CFQCSIGTYSEKEKSNSCEICPNNQITSTIGSTKKEDCFCMPGFKPASLRDETCVQCASTEWCYRVSPNSGGFWN
VASYCYSVDLSKYSKDSTTHLLCNDMSLMSSKTANFQIPCFGSQCESLYRNNEVISNNNINSLMVVRCKEGSSGF
LCDDCEDGYAKIDGLLGEVGSCKKCSLLNFTPFILSNLFIFGLIIYSVWVLRLEPTANEEEMPIVQVTIIRTTIQ
HMQFLGLLITTNVVRFEYFPNLTVAISYLGSSNSFLPLFQCLLSKFGIDRHGPEALKIYSIVISTMPFLLIIFAI
IAAPILRYFREKEVRGFLLGRQEMEGECLQFQKSYFSWVFSLFIVSFFCYFSMSIKNIYSIISCTSLNIDIQNPT
NSLMGSYVTPESMNLFQNTNNEIPGLLSVVSIAKENICWSHGHFVAIFYSLIGLVFWHILIPIVYLLCIYFDTKN
DKIQRRRLVYGWWISGYENTNMNIWEITTYLSQIIYLIVILLIGILHTFNYVVAYYAVPGGSHYETLNPLCSPIL
SSFCCIVLTISLDSIYSFVRPNSQDFLRTGIQKVDISSVGVEDELLTEEQKQKQKVAYGGVNSKQTSRLNWSSYF
YFIRRMSSFSIITVIFASILPRLNVNIENKIENLRGTWIDSVDTSLIPLGYGDFSIRFFSIFSVIVNYFFIFIIL

FSVLGIKIYKRVLPMLSSKSKVEVNIKSMSDFASSSDSIKQVNKGNQVNGRISNIGNAFVSYHDMNKTSQIGDQD
GERVDDEISIEQLIRMHLRTDCLGFLSHEDRVSVVVAIKRCDNRQEGLAISLRATLDSRFGESGKYTLIENNLFV
CESLRSYGFSD

>CPATCC_001478 [Cryptosporidium parvum Iowa II] XP_625616.1
CTPCPENTVGLTTNLYSCVNCPINSSTYGDTGLYSLDSCICLPGFFKLESTNSCISCPLNSQKYVCPGKFDSNYC
TTNEKLKNGGWWFSDNYVGIVIKCPHEEACLPECKGCKEGYTGPLCNTCTSGYFPLEYGLDKSCIECPNQYLLLV
AFIGMIMLVLLISYGIARVSELSIKYTALSRRNSAVIAKFKNLLFFISISTTYGIFVKYKDSEFQNMDEFFIHYL
KIIKLLSPNLWLPLGIFPSKCLFKLFPIIMQSVGQVNLTDANADLFIANSVPILGIAILFITFTIKHIIDVNKIY
KENDTKTELNRMKVMESSARLEKFNKILKDQTEKSTYRFICTSQVYLTLIYSTVSWSIIVSMTCTNIPGFGTVQS
QCYLLPCKTLNSDALIGYIYYCTMFPLINIISGIVIRKKIIRQNQLYTISSSFLFSGYKISVLYWESVRFYIISG
IINVLNLNYVFARYLILFVLSIYSILVFFESIISPVDDSDSSKKNGMDKPWWIFSTSLNINTIISGFTLIFIPCL
LCFYSYYPKFEFIIEITIMTFHIGTAFFMIQQVFVDTFFLIHQYNKHKQLYGDKEQILDAKESNDGLSLYYKHKL
GDNEQTNFMLGLLKSPFPEEAAAQMLYLIEQGNIFDCEAKDIGKTLYSAGFYEPQGQTLRYLIPKIKGYRSSSDI
PVENETELLGEVRKCVMNTIRILGIESAVKNSQFRAKGARVIFSSQQFGESTLLKTASDDLFKFLSDIRCSNSKE
